# Inducible and reversible SOD2 knockdown in mouse skeletal muscle drives impaired pyruvate oxidation and reduced metabolic flexibility

**DOI:** 10.1101/2024.09.23.614547

**Authors:** Ethan L. Ostrom, Rudy Stuppard, Aurora Mattson-Hughes, David J. Marcinek

## Abstract

Graphical Abstract

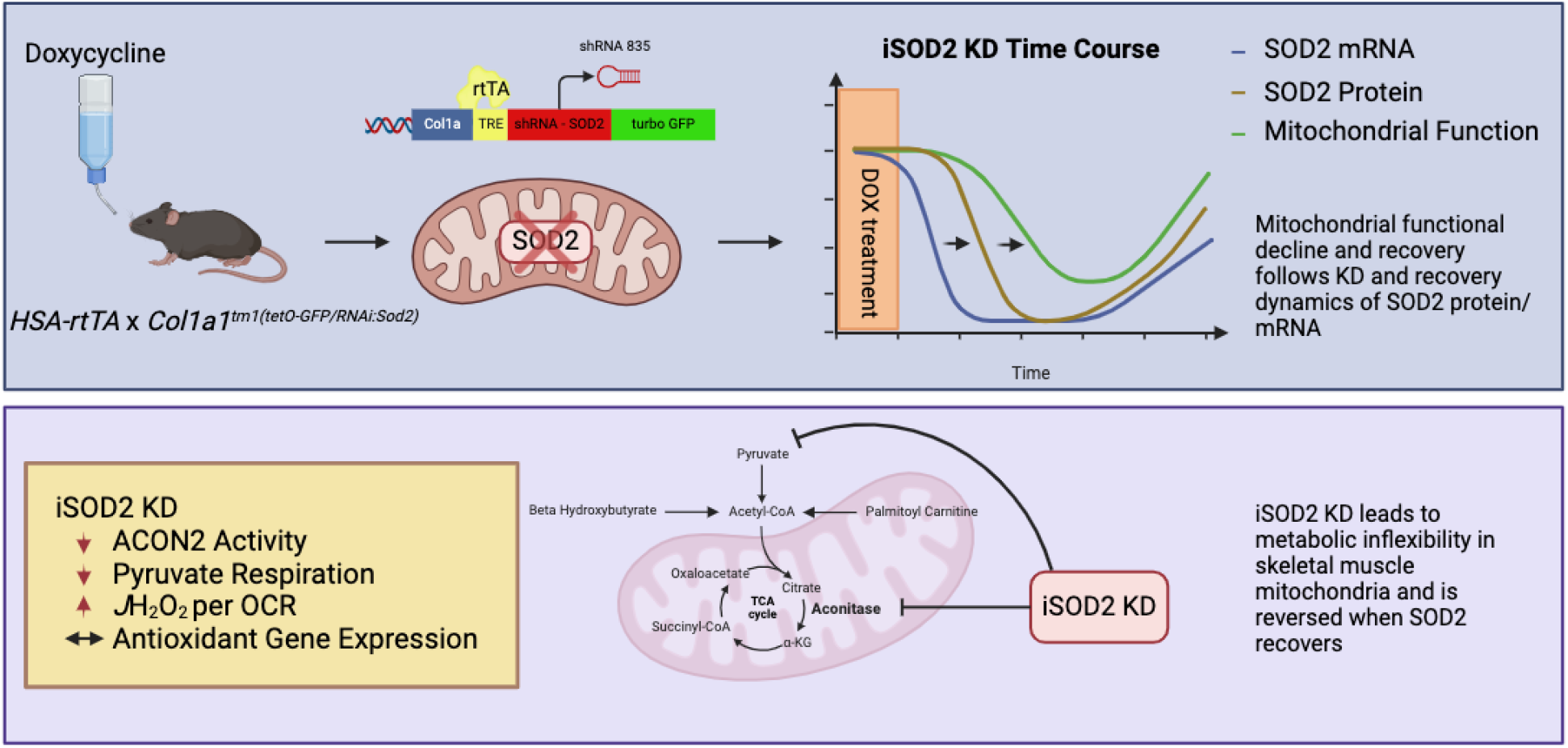

**Highlights:**

- SOD2 knockdown and recovery is achieved in skeletal muscle by using a shRNA targeted to SOD2 mRNA controlled by a tetracycline Response Element and reverse tetracycline transactivator protein
- SOD2 KD is induced by administering doxycycline in the drinking water
- Mitochondrial functional decline and recovery follows the time course of SOD2 protein decline and recovery
- Sustained SOD2 KD precipitates reduced metabolic flexibility in skeletal muscle mitochondria characterized by impaired pyruvate respiration in the presence of other substrates

**Introduction:** Skeletal muscle mitochondrial dysfunction is a key characteristic of aging muscle and contributes to age related diseases such as sarcopenia, frailty, and type 2 diabetes. Mitochondrial oxidative distress has been implicated as a driving factor in these age-related diseases, however whether it is a cause, or a consequence of mitochondrial dysfunction remains to be determined. The development of more flexible genetic models is an important tool to test the mechanistic role of mitochondrial oxidative stress on skeletal muscle metabolic dysfunction. We characterize a new model of inducible and reversible mitochondrial redox stress using a tetracycline controlled skeletal muscle specific short hairpin RNA targeted to superoxide dismutase 2 (iSOD2).

**Methods:** iSOD2 KD and control (CON) animals were administered doxycycline for 3-or 12-weeks and followed for up to 24 weeks and mitochondrial respiration and muscle contraction were measured to define the time course of SOD2 KD and muscle functional changes and recovery.

**Results:** Maximum knockdown of SOD2 protein occurred by 6 weeks and recovered by 24 weeks after DOX treatment. Mitochondrial aconitase activity and maximum mitochondrial respiration declined in KD muscle by 12 weeks and recovered by 24 weeks. There were minimal changes in gene expression between KD and CON muscle. Twelve-week KD showed a small, but significant decrease in muscle fatigue resistance. The primary phenotype was reduced metabolic flexibility characterized by impaired pyruvate driven respiration when other substrates are present. The pyruvate dehydrogenase kinase inhibitor dichloroacetate partially restored pyruvate driven respiration, while the thiol reductant DTT did not.

**Conclusion:** We use a model of inducible and reversible skeletal muscle SOD2 knockdown to demonstrate that elevated matrix superoxide reversibly impairs mitochondrial substrate flexibility characterized by impaired pyruvate oxidation. Despite the bioenergetic effect, the limited change in gene expression suggests that the elevated redox stress in this model is confined to the mitochondrial matrix.

## Introduction

Skeletal muscle mitochondria play a critical role in maintaining muscle function and muscle and systemic metabolic health. Mitochondria respond to periods of fasting, feeding, and exercise in a coordinated fashion to maintain energy and redox homeostasis. Aging and diseases of skeletal muscle are consistently associated with mitochondrial dysfunction characterized by declining P/O ratio, increased reactive oxygen species (ROS) production, impaired ADP sensitivity, and impaired fuel utilization (Marcinek, Schenkman et al. 2005, Siegel, Kruse et al. 2011, Campbell, Duan et al. 2019, Pharaoh, Brown et al. 2021, Campbell, Djukovic et al. 2023). Oxidative distress occurs when ROS production exceeds the capacity of the antioxidant defenses and has been implicated in the etiology of many diseases of skeletal muscle, including sarcopenia, muscle weakness, insulin resistance, and diabetes (Anderson, Lustig et al. 2009, Jang, Lustgarten et al. 2010, Andersson, Betzenhauser et al. 2011, Fisher-Wellman and Neufer 2012). The major ROS species produced in muscle is superoxide (Brand 2016). Under basal resting conditions in skeletal muscle, superoxide is primarily generated from mitochondria particularly in conditions of excess nutrient availability and low ATP demand (excessive calorie consumption and sedentary behavior). These conditions create a high NADH:NAD+ ratio and high electrochemical potential gradient causing backpressure against matrix dehydrogenases and the electron transport system which drive superoxide production (Fisher-Wellman and Neufer 2012, Fisher-Wellman, Gilliam et al. 2013, Fisher-Wellman, Lin et al. 2015, Goncalves, Quinlan et al. 2015). Superoxide can react directly with iron sulfur clusters or react to form other more reactive radical species such as peroxynitrite (ONOO^−^) or hydroxyl radical (•OH) which drive macromolecule oxidation and oxidative distress (Sies, Belousov et al. 2022). Although elevated mitochondrial ROS production and altered redox homeostasis are consistent features associated with mitochondrial dysfunction, it is still debated whether oxidative distress itself causes metabolic dysfunction or if it is simply correlated with age-related declines. To better understand the role of mitochondrial oxidative distress on skeletal muscle metabolic dysfunction more robust models of mitochondrial oxidative distress are needed.

Genetic manipulation of tissue specific and compartment specific antioxidant enzymes are the most reliable and offer the cleanest way to alter redox status *in vivo*. Superoxide dismutase 2 (SOD2) is localized to the mitochondrial matrix and catalyzes the conversion of superoxide to hydrogen peroxide, a critical enzyme in the antioxidant defense system (Sies, Belousov et al. 2022). However there remain limitations to traditional approaches of genetic knockouts. Constitutive knockout of skeletal muscle SOD2 from birth display marked mitochondrial and skeletal muscle function impairments (Ahn, Ranjit et al. 2019), but it is difficult to tease apart developmental effects versus the effects of the knockout in mature adult tissue. This is an important distinction because the onset of mitochondrial redox distress associated with aging and disease does not manifest until after the organism is fully developed. Inducibility circumvents this problem of developmental effects as demonstrated by inducible models showing less severe phenotypes than earlier constitutive models (Zhuang, Yang et al. 2021). Furthermore, the ability to reverse the knockdown would provide another layer of control to test adaptive and pathogenic responses to mitochondrial redox stress more directly (Cox, McKay et al. 2018). This is an important scientific control, allowing researchers to test whether elevations in mitochondrial superoxide levels are necessary and sufficient to induce phenotypic changes in mitochondrial and muscle function.

The goal of this project was to characterize a model of inducible and reversible skeletal muscle SOD2 Knockdown (iSOD2 KD) adapted from Cox et al.(Cox, McKay et al. 2018). Here we target SOD2 KD to striated muscle by crossing the iSOD2 KD mice with the Human Skeletal Actin – rtTA mice (Iwata, Englund et al. 2018) to better understand the metabolic and physiological responses to elevated steady state mitochondrial superoxide levels in adult muscle. This model uses RNA interference to silence the SOD2 mRNA in skeletal muscle to prevent SOD2 protein expression. We measured mitochondrial and muscle function in response to SOD2 knockdown and recovery in mature male and female mice to test the effect of mitochondrial oxidative distress on respiration and contractile function. We demonstrate that SOD2 KD causes impaired pyruvate driven respiration in permeabilized muscle fibers in the presence of other respiratory substrates. The SOD2 KD also leads to elevated ROS production per unit of oxygen consumption, impairs mitochondrial aconitase activity, and increases skeletal muscle fatiguability. Importantly, the bioenergetic changes in mitochondria are reversible with recovery of SOD2 protein indicating that increases in superoxide steady state levels in mitochondria are sufficient to drive changes in mitochondrial energetics. This model will be beneficial for understanding the role of elevated steady state superoxide levels on mitochondrial bioenergetics and skeletal muscle function.

## Methods

### Animal breeding

The inducible Superoxide Dismutase 2 knockdown (iSOD2 KD) animals (Cox, McKay et al. 2018) were purchased from Jackson Labs (B6.Cg-*Col1a1^tm1(tetO-GFP/RNAi:Sod2)Gsha^*/J, strain #: 032649) and crossed with the Human Skeletal Actin driven reverse-tetracycline transactivator mouse (HSA-rtTA mouse) previously described by (Iwata, Englund et al. 2018) after genotypes were confirmed. Pups from all cages were weaned at 3 weeks of age and genotyped by tail snip and ear punch biopsies described in detail below. HSA-rtTA^+^ / iSOD2^+^ animals (double positive) were considered experimental animals, while HSA-rtTA^+^ / iSOD2^−^ animals were used as littermate controls (Supplementary Figure 1). This ensures both groups can receive DOX administration, but only the double positive experimental animals will experience SOD2 KD. This provides control for any potential confounding effects of rtTA expression and its interaction with DOX on mitochondrial and muscle physiology in the absence of SOD2 KD. The cages that did not produce experimental animals were discontinued in the breeding scheme and euthanized per University of Washington IACUC protocols upon establishing working genetic crosses.

### Genotyping

Tail snips and ear punch biopsies were taken for genotyping during weaning. Tissue samples were digested in 10mM Tris, 50mM NaCl, 1mM EDTA, 0.5% SDS (pH 8.0) supplemented with 1% Proteinase K and incubated at 55°C overnight. Following digestion, DNA was isolated by adding 450µl of a 1:1 phenol chloroform solution, vortexing for 10 seconds followed by centrifugation at maximum RPM for 10 minutes in a tabletop microcentrifuge. The top layer of solution was aliquoted into a new tube with 1mL of 95% ethanol. Samples were vortexed and centrifuged at maximum RPMs for 10 minutes. Excess EtOH was poured off and remaining EtOH was evaporated in a Speed Vac centrifuge under high heat for 25 minutes. Following EtOH evaporation, samples were reconstituted with 75µl pure DNase/RNase free water. DNA quantity and sample purity were assessed by 260/280nm and 260/230 ratio and ng/µl using a NanoDrop spectrophotometer (ThermoScientific). Samples were diluted to a concentration of 100ng/µl in RNase/DNase free water. Five hundred nanograms of DNA were used in the PCR reaction to amplify genes of interest. Following amplification, DNA was separated by gel electrophoresis on a 1% agarose gel for 30 minutes at 140V. Gels were imaged under UV transillumination in a ChemiDoc XRS (Bio-Rad, Hercules, CA). The wild type allele for the B6.Cg-*Col1a1^tm1(tetO-GFP/RNAi:Sod2)Gsha^*/J line produces a gene product at 200 bp, and the mutant transgene produces a gene product at 267bp. The HSA-rtTA transgene produces a gene product of 747bp. Supplementary Figure 1 illustrates PCR products visualized by agarose gel electrophoresis of one experimental animal and one littermate control animal. All animals were screened for each gene to determine whether animals were experimental or littermate controls. Any animals with unwanted genotypes were euthanized per University of Washington IACUC guidelines.

### Doxycycline Administration

Animals were treated with 0.5mg/ml DOX administered in the drinking water (Doxycycline Hyclate, Sigma Aldrich St. Louis, MO, USA) supplemented with 2% sucrose for various treatment durations described in further detail below (Iwata, Englund et al. 2018).

### Model Mechanism

This model uses RNA interference controlled by an inducible TET-ON system (See Figure 1A). In TET-ON systems the reverse tetracycline transactivator (rtTA) protein is expressed but inactive. Doxycycline administration activates the rtTA protein, driving expression of the tetracycline response element (TRE) promoter and downstream genes. In this case the TRE drives expression of a short hairpin RNA that targets SOD2 mRNA for degradation and subsequent ablation of SOD2 protein (Cox, McKay et al. 2018). This model also co-expresses Turbo Green Fluorescent Protein (GFP) as an indicator of TRE promoter activation. DOX administration knocks down SOD2 protein and increases GFP fluorescence, while removal of doxycycline will restore SOD2 protein levels and decrease GFP fluorescence.

**Figure 1.**
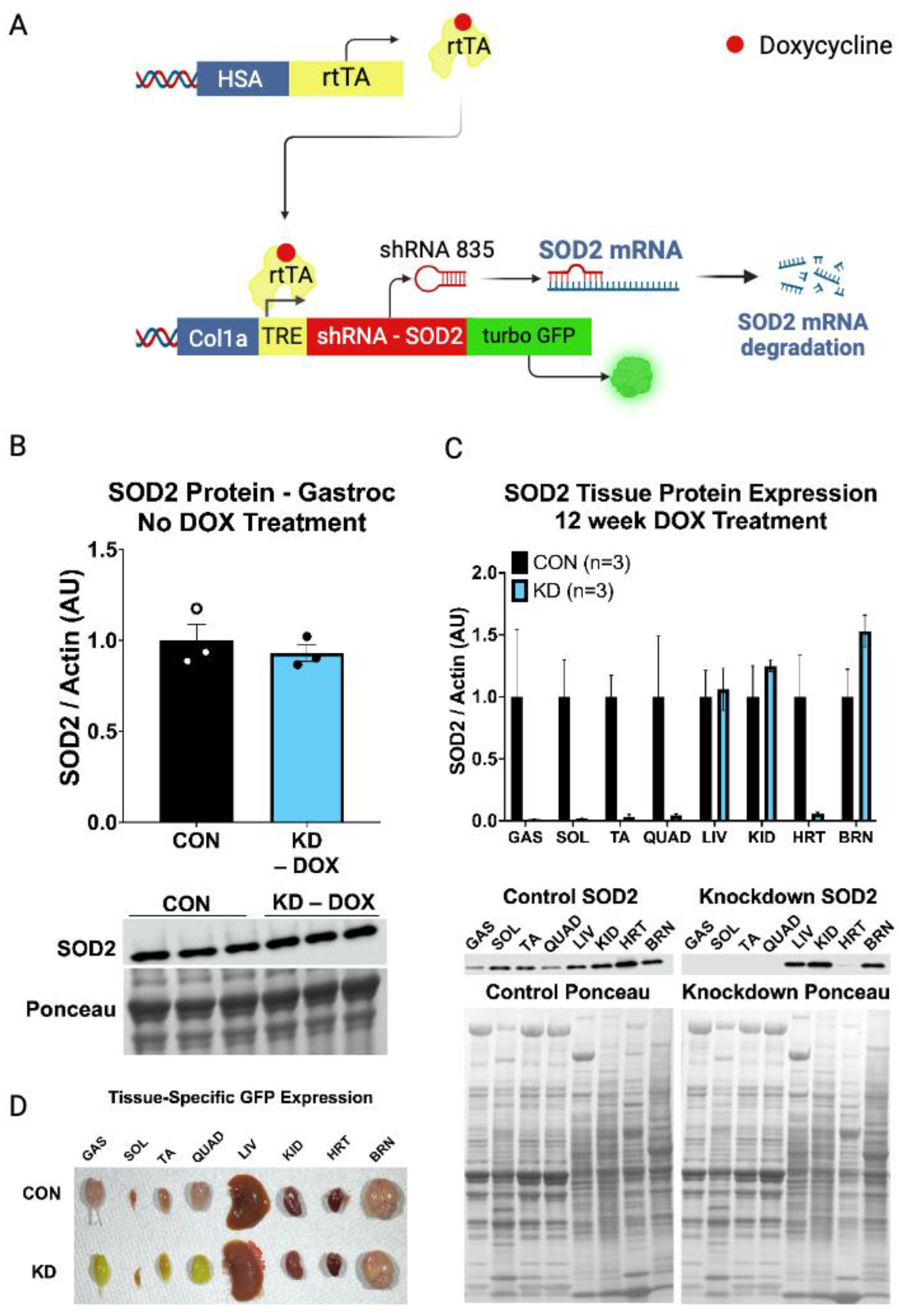
A) Inducible and reversible SOD2 knockdown is achieved with a tetracycline response element (TRE) controlling expression of a short hairpin RNA (shRNA 835) which targets SOD2 mRNA for degradation. The reverse tetracycline transactivator transcription factor is expressed only in striated muscle via the human skeletal actin (HSA) promoter. Experimental animals express both the TRE-shRNA-GFP and HSA-rtTA genes, while littermate controls only express HSA-rtTA. When experimental animals are fed doxycycline (DOX) this induces shRNA expression in striated muscle. Turbo GFP is co-expressed with the shRNA as an indicator of TRE promoter activation. B) SOD2 protein expression in gastrocnemius muscle of untreated KD (KD - DOX) and control animals to test for basal activation of the TRE promoter in the absence of DOX. C) Tissue specific expression of SOD2 in response to 12 weeks of continuous doxycycline administration in control or iSOD2 KD animals. Bands were normalized to total protein with a ponceau stain, then iSOD2 KD animals were expressed relative to the controls within the same tissue. All data presented are mean ± SD. D) Tissue-specific GFP expression is visible to the naked eye after 12 weeks of continuous DOX administration in CON and KD in skeletal muscle tissue. GAS – gastrocnemius, SOL – soleus, TA – tibialis anterior, QUAD – quadriceps, LIV – liver, KID – kidney, HRT – heart, BRN – brain. Representative image of respective tissues and western blot image of control and iSOD2 KD animals (n=3 animals per genotype). Heart muscle also shows KD of SOD2 in response to DOX administration, likely due to HSA-rtTA promoter being expressed in heart tissue. All data presented are mean ± SD.

### Study Design

To understand the physiological effects of iSOD2 KD in skeletal muscle we performed several physiological and biochemical analyses at multiple time points throughout the KD and recovery phase in young adult male and female mice. We used two different treatment conditions to induce SOD2 KD: a three-week DOX treatment and up to 24-week recovery to assess KD and recovery dynamics, as well as a continuous 12-week DOX treatment to establish sustained SOD2 KD. The effects of SOD2 KD on muscle and mitochondrial physiology were assayed at multiple time points throughout each of these treatment lengths. The timeline for each treatment phase and recovery time course is illustrated in Figure 2. All experiments used littermate controls fed doxycycline for the same time frame as iSOD2 KD’s to account for any confounding effects of doxycycline on mitochondrial and muscle physiology. There were no differences in SOD2 protein expression or GFP fluorescence between untreated KD animals and littermate controls suggesting there was no basal activation of the promoter in the absence of DOX.

**Figure 2.**
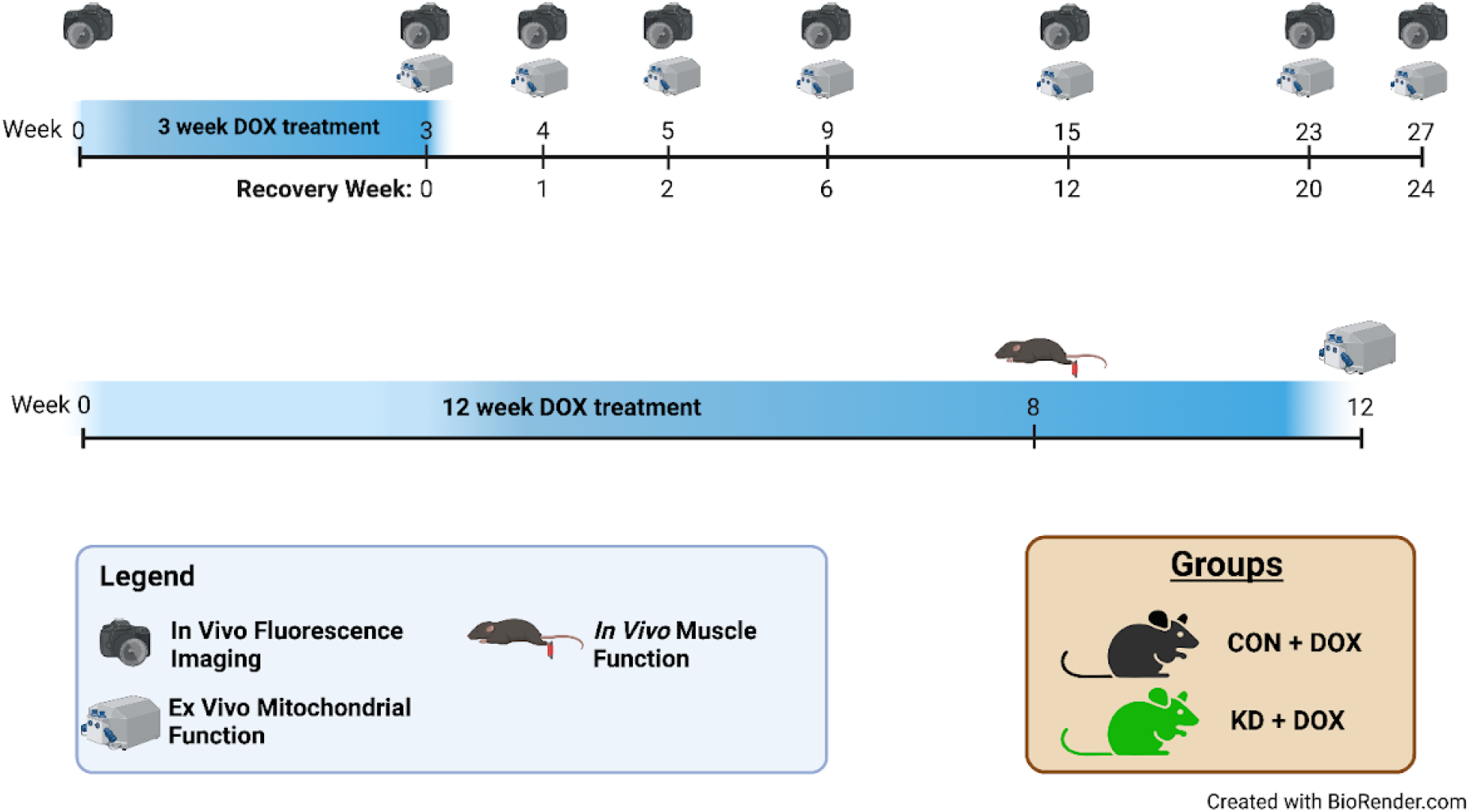
A) In the first set of experiments animals were administered DOX for 3 weeks and followed for up to six months to assess kinetics of SOD2 knockdown and recovery and the mitochondrial phenotype. Outcome measures include *in vivo* GFP fluorescence, *ex vivo* high resolution respirometry, gene expression and protein content of target genes, and mitochondrial aconitase activity as a marker of mitochondrial oxidative stress. A second set of experiments were conducted with a 12-week continuous DOX administration to follow up on the initial findings from the time course experiments. At week 8 of continuous DOX treatment a subgroup of animals were subjected to *in vivo* muscle function and fatigue testing. Animals were allowed to recover and then used for high resolution respirometry experiments. All experiments used littermate controls administered DOX for the same amount of time to control for any confounding effects of DOX on mitochondrial physiology.

### In Vivo GFP Fluorescence

GFP fluorescence was monitored before and at several time points after a 3-week doxycycline treatment using an *in vivo* IVIS Spectrum Imaging System (PerkinElmer, Waltham, MA). Briefly, animals were anesthetized with isoflurane, then the fur was removed from the right hind limb using Nair and a cotton swab. Following hair removal, the animals remained under anesthesia and were placed on their side in the IVIS Spectrum with the right hind limb up. Limbs were imaged with 465/520 (Ex/Em) lasers to visualize GFP fluorescence. The region of interest around the right tibialis anterior was selected to minimize background fluorescence from other body structures.

### In Vivo Muscle Function & Fatigue

iSOD2 KD and control animals underwent *in vivo* muscle function testing using a 1300a 3-in-1 Whole Animal System (Aurora Scientific, Aurora ON, CA). Briefly, animals were anesthetized with 4% isoflurane and maintained under anesthesia at 1 −2% on water recirculating heated pad. The animals’ right hindlimb was secured at the knee and foot taped to a footplate at 90°. Subdermal electrodes were placed behind the knee to stimulate the tibial nerve and cause plantar flexor muscle contraction. A voltage titration was used to determine optimal voltage (4-8V) followed by 2 minutes of rest. At the end of the 2-minute rest period a force frequency (FF) analysis was conducted, 1 stimulation every minute for nine minutes (10-200Hz) (Figure 5A). The maximal torque-force (mN-m) was determined from this curve. Following the FF, animals rested for 5 minutes, followed by a fatiguing protocol (100Hz frequency, 0.2ms pulse duration, every 4 seconds) for 120 contractions (Figure 5B). Animals were allowed to recover in their home cages after completion of the *in vivo* muscle function testing. Animals were maintained on DOX treatment until euthanasia and tissue collection for O2K experiments and biochemical assays.

### Ex Vivo Mitochondrial Respiration

High resolution respirometry was performed on permeabilized red gastrocnemius muscle fibers. Gastrocnemius fibers were carefully removed and placed in ice cold BIOPS followed by fiber bundle mechanical separation using fine forceps and scissors. Fiber bundles were weighed (1.5-3 mg tissue) and then permeabilized with 50µg/ml Saponin in BIOPS buffer for 40 minutes. Fibers were placed in a 2ml chamber of an Oxygraph 2K dual respirometer/ fluorometer (Oroboros Instruments, Innsbruck, Austria) at 37°C and stirred gently during substrate uncoupler and inhibitor titration (SUIT) protocols. SUIT protocols with different substrate orders were used to test the isolated and additive effects of substrate utilization on oxygen consumption, but generally included assessment of leak respiration, Complex I driven respiration, maximal oxidative phosphorylation, uncoupled respiration, Complex II respiration, and complex IV respiration. The substrates used were glutamate (10mM), beta hydroxybutyrate (25mM), palmitoyl carnitine (2mM), pyruvate (5mM), and succinate (10mM). Each substrate titration included malate (2mM) prior to stimulation with ADP (2.5mM). Substrate order as well as concentrations for each protocol used are shown in table 1.

**TABLE 1.** SUBSTRATE UNCOUPLER INHIBITOR TITRATION (SUIT) PROTOCOLS. Four substrate uncoupler inhibitor titration protocols were used to assess substrate driven respiration in KD and control muscles. Comparing the same sample across different protocols provides additional insight into substrate level control of respiration and additive effects of different substrates based on the order of addition (Step 1-10).

| STEP | Protocol 1 |  | Protocol 2 |  | Protocol 3 |  | Protocol 4 |  |
| --- | --- | --- | --- | --- | --- | --- | --- | --- |
|  | Substrate | [Final] | Substrate | [Final] | Substrate | [Final] | Substrate | [Final] |
| 1A. | Glutamate | 10 mM | Pyruvate | 5 mM | Palmitoyl Carnitine | 50µM | Beta-Hydroxybutyrate | 10mM |
| 1B. | Malate | 2 mM | Malate | 1 mM | Malate | 1 mM | Malate | 1 mM |
| 2 | ADP | 150µM | ADP | 150µM | ADP | 150µM | ADP | 150µM |
| 3 | ADP | 2.5 mM | ADP | 2.5 mM | ADP | 2.5 mM | ADP | 2.5 mM |
| 4 | Pyruvate | 5 mM | Glutamate | 10 mM | Pyruvate | 5 mM | Pyruvate | 5 mM |
| 5 | Succinate | 10 mM | Palmitoyl Carnitine | 500µM | Glutamate | 10 mM | Glutamate | 10 mM |
| 6 | FCCP | 0.5 µM | Succinate | 10 mM | Succinate | 10 mM | Succinate | 10 mM |
| 7 | Rotenone | 0.5 µM | Rotenone | 0.5 µM | Rotenone | 0.5 µM | Rotenone | 0.5 µM |
| 8 | Antimycin A | 2.5 µM | Antimycin A | 2.5 µM | Antimycin A | 2.5 µM | Antimycin A | 2.5 µM |
| 9A. | Ascorbate | 2mM | Ascorbate | 2mM | Ascorbate | 2mM | Ascorbate | 2mM |
| 9B. | TMPD | 500µM | TMPD | 500µM | TMPD | 500µM | TMPD | 500µM |
| 10 | AZD | 100mM | AZD | 100mM | AZD | 100mM | AZD | 100mM |

### Citrate Synthase Activity

Citrate synthase catalyzes the conversion of oxaloacetate and acetyl-CoA to citrate and is a marker of mitochondrial volume and density in skeletal muscle (Larsen, Nielsen et al. 2012). CS activity was assessed using a kit from Sigma Aldrich (Catalog No. MAK193-1KT Sigma Aldrich St. Louis, MO, USA). Briefly, whole gastrocnemius tissue lysate was homogenized with Cell Lytic-MT buffer supplemented with 0.1% protease and 0.2% phosphatase inhibitors and spun down for 5 minutes at 10,000xg. CS positive controls or sample were loaded in designated wells of a 96 well plate in duplicate, along with complete assay buffer consisting of buffer, acetyl-CoA, and DTNB. Background was determined using a kinetic read at 412 nm prior to adding oxaloacetate (OAA). To initiate the forward reaction, OAA was quickly added using a multichannel pipet followed by kinetic reads at 412 nm. CS rate of activity was determined by subtracting background at each timepoint followed by averaging duplicates and correcting for protein loading.

### Aconitase Activity Assay

Mitochondrial aconitase (ACON2) catalyzes the reaction of citrate to cis-aconitate in the TCA cycle. Mitochondrial aconitase is considered a good marker of mitochondrial oxidative stress because the active site of the enzyme is redox sensitive (Li, Pedersen et al. 2001, Figueiredo, Powers et al. 2009). Here we use the aconitase activity assay (Catalog No. ab109712, Abcam, Cambridge, UK) to assess the extent of mitochondrial oxidative stress triggered by SOD2 KD. Mitochondria were isolated from quadriceps muscles of KD or CON animals treated with DOX. Briefly, tissue was homogenized with a tight-fitting Dounce homogenizer in ice cold mitochondrial isolation buffer (210mM Sucrose, 2mM EGTA, 40mM NaCl, 30mM HEPES, pH 7.4). After homogenization samples were spun 900xg for 10 minutes at 4°C. Following centrifugation, the supernatant was aliquoted into a new 1.5mL tube. Samples were then spun again at 10,000xg for 10 minutes at 4°C to pellet mitochondria. Supernatant was discarded and the mitochondrial pellet was resuspended in 10mM Tris, 1mM EDTA. A Bradford assay was used to determine mitochondrial protein content. On the day of the assay, mitochondria were diluted to 0.5µg/µl with aconitase preservation solution (Abcam) and 50 µl were loaded in duplicate to designated wells. Four CON samples were treated with 100µM H_2_O_2_ in addition to the standardized assay conditions as a negative assay control. After the addition of assay buffer and isocitrate, a kinetic read was used at 240 nm every minute for 30 minutes to determine mitochondrial aconitase activity.

### Electron Transport System Complex Activities

The electron transport system (ETS) activity protocols were adapted from several previously published methods (Janssen, Smeitink et al. 2003, Luo, Long et al. 2008, Barrientos, Fontanesi et al. 2009, Medja, Allouche et al. 2009, Chen, Thorburn et al. 2011, Spinazzi, Casarin et al. 2011, Spinazzi, Casarin et al. 2012). Briefly, the mouse gastrocnemius tissue was manually disrupted using a Dounce (glass/glass) homogenizer prior to gentle centrifugation at 660 g to remove debris. Mitochondria were permeabilized by repeated (N = 3) freeze thaw cycles. ETS complex activities were measured using UV Vis spectrophotometry to detect changes in absorption of reporter compounds as electrons were donated or accepted. Data were collected using a Cary 60 spectrophotometer (Agilent, Santa Clara, CA). All ETS complex rates were determined as the difference of activity measured in the absence and presence of complex-specific inhibitors and normalized to the total protein included in each reaction.

### RT-qPCR

RNA was isolated from pulverized gastrocnemius muscle using the RNeasy Fibrous Tissue Mini Kit (Qiagen, Germany, Catalog No. 74704). RNA quality and quantity was assessed using a Nanodrop spectrophotometer (ThermoScientific, USA). All samples had 260/280 and 260/230 ratios > 1.8. For target gene amplification, exon-spanning primers were used from Themo Scientific. A full list of primers is available in the supplementary information (Suppl. Table 2). Verso 1-Step RT - qPCR kit (Themo Scientific, USA catalog no. AB4101C) was used per the manufacturer’s instructions for cRNA library preparation and qPCR amplification. Differential gene expression was analyzed using the ΔΔCt method with RPL41 as an internal housekeeping gene. PCR was run on an Applied Biosystems 7900 HT RT–qPCR machine compatible with 384-well plates. All samples were run in duplicate.

### Western Blotting

Tissues were cryo-pulverized, and cells were homogenized and lysed with Cell Lytic MT lysis buffer supplemented with 0.1% protease inhibitors, 0.2% phosphatase inhibitors, followed by centrifugation at 1000 xg and 4°C for 5 minutes. After protein concentrations were measured using the Bradford protein assay, samples were diluted to 2µg/µl with lysis buffer. Twenty micrograms of protein were loaded per well for each western blot, followed by SDS-PAGE and semi-dry transfer to a nitrocellulose membrane. Protein targets were normalized to total protein loaded per lane using a Ponceau S stain. Following de-staining, membranes were blocked with blocking buffer then incubated with primary antibody overnight at 4°C and with secondary antibodies for 1 hour at room temperature followed by imaging. A list of primary antibodies, the manufacturer, their dilutions, and diluent recipes are shown in Supplementary Table 1.

### Statistical Analysis

Statistical analysis was performed using GraphPad Prism (La Jolla, CA) and SPSS software (IBM, Armonk NY). Different statistical tests were used depending on the outcome metric and grouping factors. Independent sample t-tests were used when comparing strict group differences (KD vs CON). When including both sexes, a 2 x 2 ANOVA (Genotype x Sex) was used. For post hoc multiple comparison testing, Tukey’s Test was used. For force frequency analysis Group x Sex x Frequency ANOVA (2 x 2 x 9) was performed with repeated measures on the Frequency factor. For fatigue analysis, the shape of the curves were compared using a LOESS model to fit the data and area under the curve was calculated to determine fatigue resistance between genotypes.

## Results

### SOD2 knockdown and recovery dynamics in response to three weeks of doxycycline treatment

We sought to characterize the temporal dynamics of SOD2 protein levels, SOD2 mRNA, and GFP fluorescence in skeletal muscle of iSOD2 KD and control animals before and following a three-week DOX administration (Figure 2). Since this model uses an RNA interference mechanism, the time course of protein KD is dependent on the half-life of SOD2 protein at the start of DOX treatment and is therefore delayed relative to the decline of SOD2 mRNA. Subgroups of animals were euthanized at each time point to collect tissues for SOD2 gene expression and protein content in skeletal muscle as well as enzyme activity assays and high resolution respirometry measurements. SOD2 protein content was not different between iSOD2 KD and controls in the absence of doxycycline demonstrating no basal activation of the TRE promoter (Figure 1B). This model successfully knocked down SOD2 in skeletal muscles, in gastrocnemius, soleus, tibialis anterior, and quadriceps muscles, as well as heart tissue, without changes in liver, kidney, or brain (Figure 1C). GFP is easily visualized in KD skeletal muscle, but not other tissues compared to controls after 12 weeks of continuous DOX administration (Figure 1D).

There were no differences in GFP fluorescence prior to starting the DOX treatment but after three weeks of DOX administration there was a significant increase in *in vivo* GFP fluorescence in KD but not controls (Figure 3A). GFP fluorescence decreased dramatically by 6 weeks of recovery but remained elevated above control levels until 24 weeks of recovery. SOD2 mRNA showed significant knockdown after 3 weeks of DOX treatment and remained undetectable until 20 weeks of recovery and was partially recovered at 24 weeks of recovery (Figure 3B). As expected, the dynamics of SOD2 protein expression was delayed relative to the mRNA, showing a 50% reduction in protein content at 3 weeks and became undetectable by the 6th week after stopping DOX (Figure 3C). The time course of SOD2 protein knockdown is in line with previous work from our lab demonstrating SOD2 has a half-life of approximately 20 days in adult mouse skeletal muscle (Kruse, Karunadharma et al. 2016). SOD2 protein showed the same recovery dynamics as mRNA with partial recovery at 20 and 24 weeks.

**Figure 3.**
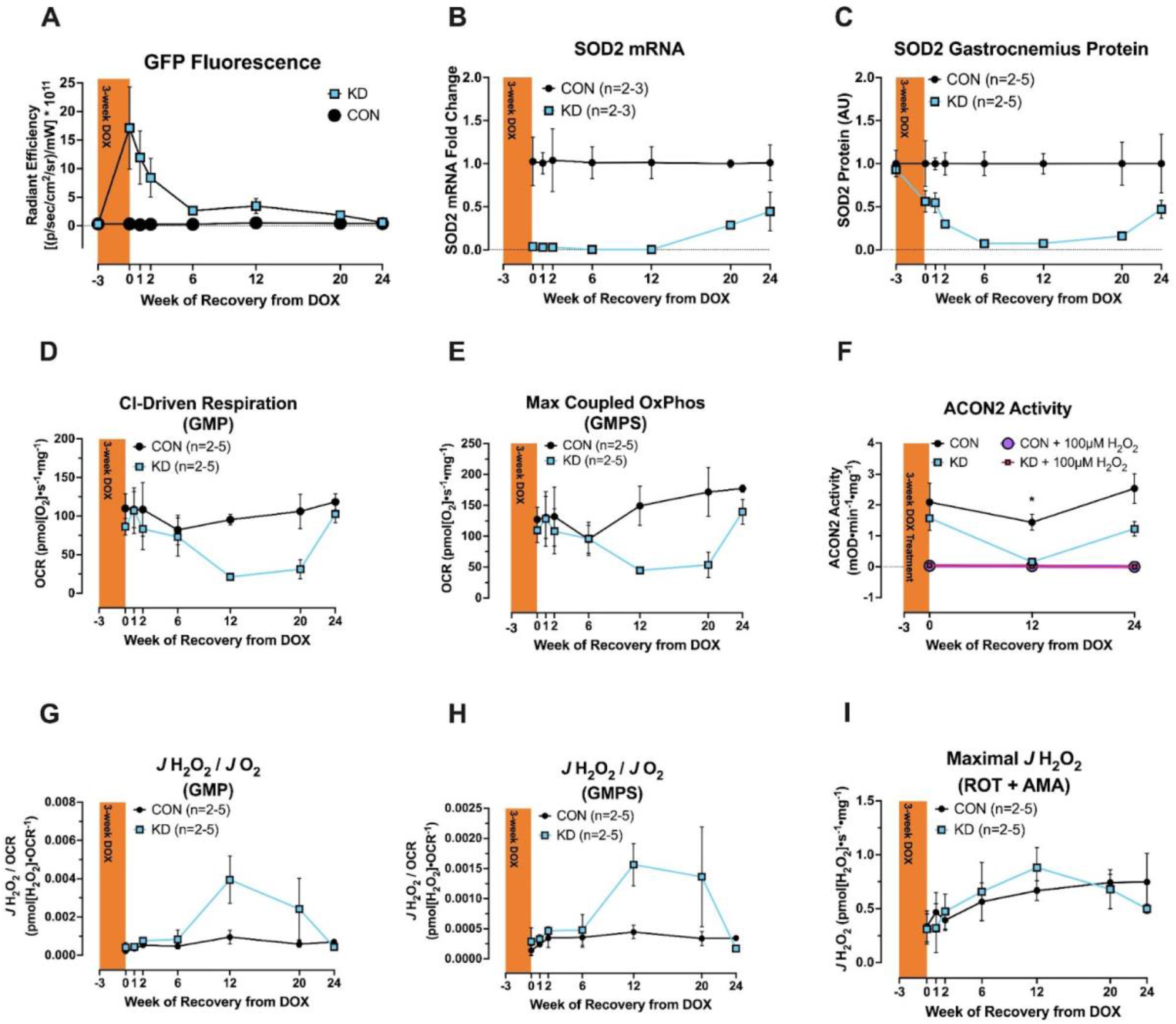
Time course of SOD2 Knockdown in response to three weeks of doxycycline (DOX) administration. A) GFP fluorescence (genotype x time interaction p<0.0001). B) SOD2 mRNA expression in gastrocnemius muscle (effect of genotype p=0.0001). C) SOD2 protein content in gastrocnemius muscle (genotype x time interaction p<0.0001). D) Complex I driven respiration in permeabilized red gastrocnemius muscle (genotype x time interaction p=0.0158). E) Maximal coupled oxidative phosphorylation (genotype x time interaction p=0.003). F) Mitochondrial aconitase (ACON2) activity at week 0, week 12, and week 24 of recovery from DOX administration (effect of genotype p=0.0004, Post hoc test at week 12, *p=0.0401). All samples show complete loss of ACON2 activity when pre-incubated with 100µM hydrogen peroxide (purple & pink lines respectively). G) ROS production rate per OCR with GMP on board (main effect of genotype p=0.0042, genotype x time interaction p=0.0002). H) ROS production rate per OCR with GMPS on board (main effect of genotype p=0.0001, genotype x time interaction p=0.0006). I) Maximal ROS production rate measured after the addition of Rotenone and Antimycin A in permeabilized muscle fibers. For the high resolution respirometry experiments, protocol 1 (Table 2) was used. Complex I driven respiration included glutamate, malate, pyruvate, and saturating ADP levels (Panel D). Maximal oxidative phosphorylation (Panel E) included glutamate, malate, pyruvate, ADP, and succinate. J H2O2 – ROS production rate, OCR – Oxygen Consumption Rate. All data presented are mean ± SD.

For the time course experiment, mitochondrial respiration and ROS production was assessed in permeabilized red gastrocnemius fibers using SUIT Protocol 1 (see Table 1 for substrate order & concentrations). SOD2 KD mice displayed impaired complex I driven respiration (Figure 3D, p<0.0001) and maximal oxidative phosphorylation capacity (Figure 3E, p<0.0001) compared to controls. This was primarily driven by differences at the 12- and 20-week timepoints. For visualization purposes, only complex I driven respiration and maximal OxPhos capacity are graphed over time. Full data sets with all substrate and inhibitor titrations are shown in Supplementary Figure 3. Mitochondrial aconitase activity (ACON2) was assessed as a marker of mitochondrial oxidative stress. ACON2 activity was significantly lower in KD compared to controls which was driven by differences at the 12-week time point (Figure 3F, main effect of genotype p=0.0004, Tukey’s post hoc comparison at 12-week recovery *p=0.0401). ROS production per rate of O_2_ consumption was significantly higher in KD compared to controls during CI driven state 3 respiration (Figure 3G), or maximal oxidative phosphorylation (Figure 3H). There was an elevation in maximal rates of ROS production in KD although this did not reach statistical significance (Figure 3I). The largest ROS production rate observed in SOD2 KD muscle occurred at week 12 corresponding to the lowest values of OCR and ACON2 activity.

### Mitochondrial biogenesis and antioxidant genes show minimal changes in response to SOD2 KD and recovery

Since mitochondrial ROS production is known to have retrograde signaling effects, we measured gene expression responses of antioxidant and mitochondrial genes at week 0, week 12, and week 24 of recovery. There were no main effects of genotype for any gene, or any genotype effects at a specific time point. There was significant genotype by time interaction effects for GRX2 (Figure 4F, p=0.0165), and PRDX3 (Figure 4H, p=0.0489) which encode mitochondrial matrix antioxidant proteins responsible for protein glutathionylation and reducing hydrogen peroxides, respectively. Despite significant interaction effects the SOD2 KD decreases antioxidant gene expression at week 12 of recovery, opposite of what would be expected for an adaptive response to elevated superoxide production.

**Figure 4.**
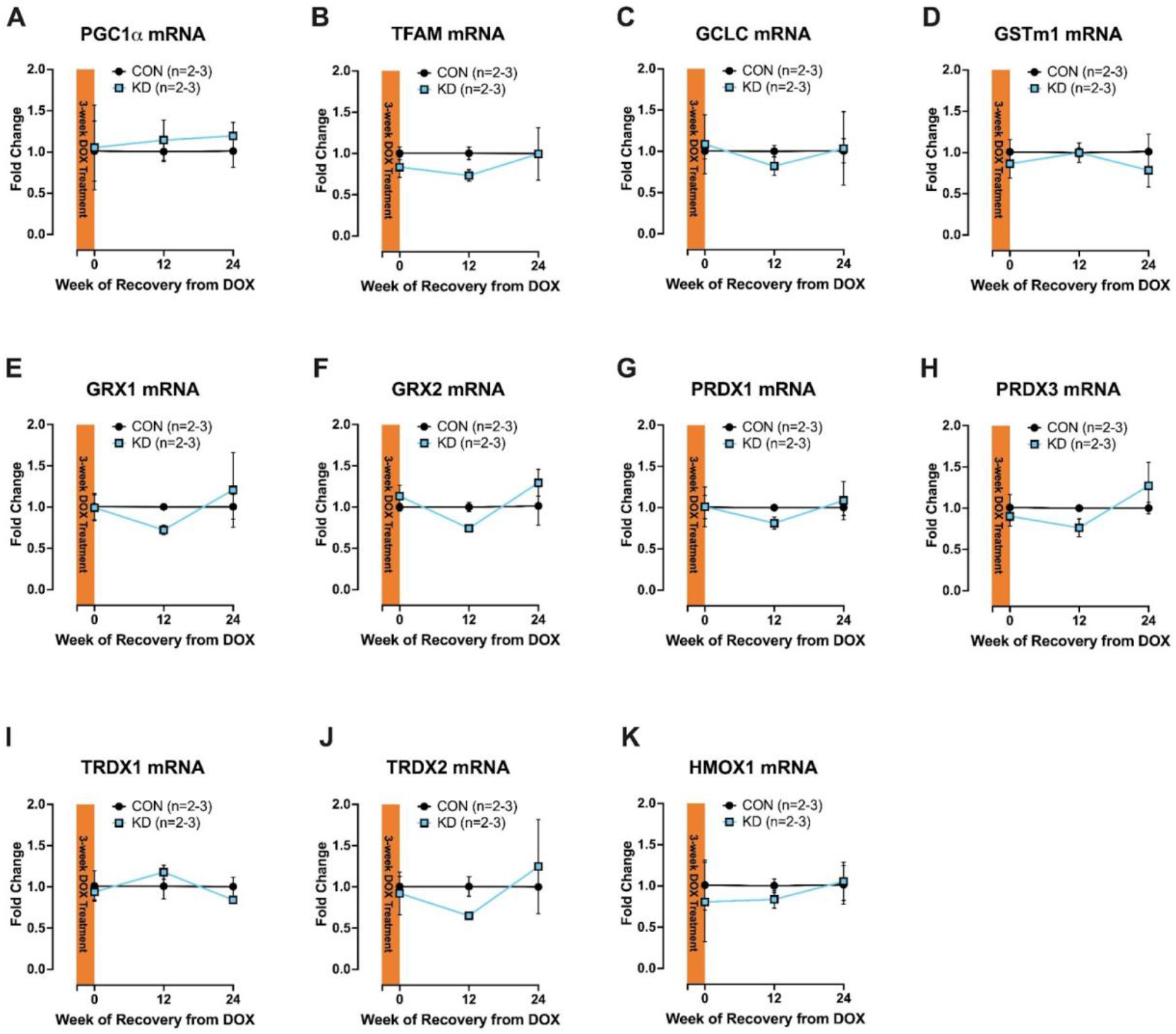
Gene expression of select antioxidant and mitochondrial biogenesis genes at recovery week 0, 12, and 24 following 3 weeks of DOX administration. A 2×3 Repeated measures ANOVA was used to test for effects of genotype, time and the genotype x time interaction. F) GRX2 showed a significant genotype by time interaction (p=0.0165). H) PRDX3 showed a significant genotype by time interaction (p=0.0489). There were no significant effects at individual time points between genotypes. RPL41 was used as a housekeeping gene. All data presented are mean ± SD.

### iSOD2 KD shows changes in skeletal muscle fatigue at 12 weeks of KD

After establishing a timeline of SOD2 KD, resulting in mitochondrial functional decline and recovery, we wanted to determine whether sustained mitochondrial redox stress affected *in vivo* skeletal muscle function. For these experiments male and female mice were treated with DOX continuously for 12 weeks. At week eight of DOX treatment *in vivo* muscle force and fatigue was assessed. There were no significant effects of genotype within each sex in force frequency metrics, so males and females were combined for subsequent analysis (see supplementary Figure S4 for data graphed by sex). A 2 x 9 repeated measures ANOVA (genotype x frequency) was used on force frequency. There were no significant main effects of genotype or genotype by frequency interactions (Figure 5A). Following force frequency testing, muscles were exposed to 100Hz isometric contractions every 4 seconds for 120 contractions to assess muscle fatigue. Although the overall shape of the curves was similar, the iSOD2 KD muscle showed reduced fatigue resistance measured by area under the curve (Figure 5B inset, independent sample t-test, p=0.0359). Animal body masses increased over the 12-week experiment, but this was not different between groups, and can be explained by the normal maturation process (Supplementary Figure S4).

**Figure 5.**
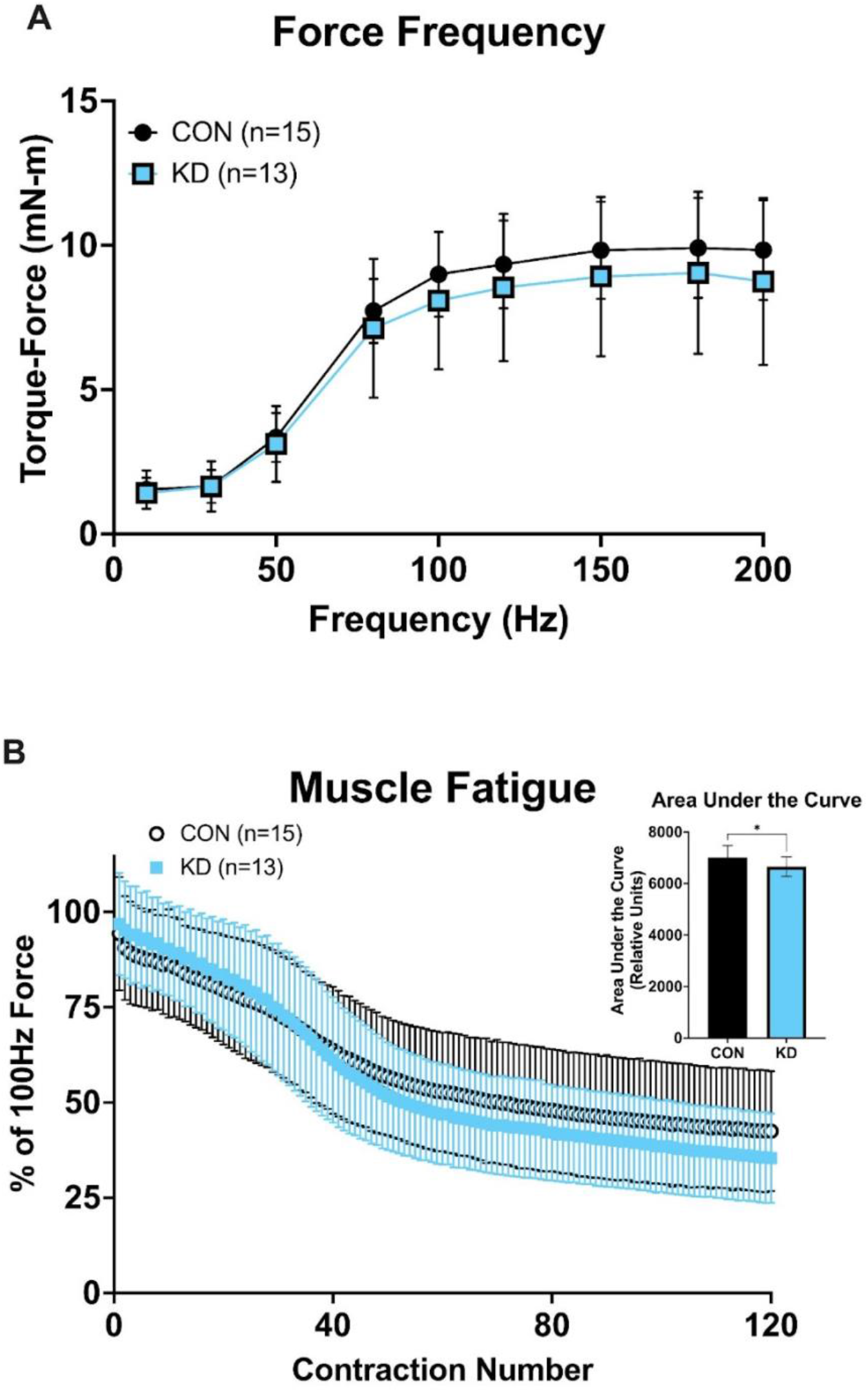
*In vivo* skeletal muscle function and fatigue following 8 weeks of continuous DOX administration. A) Force frequency curve. B) Maximal contraction rate. C) Maximal relaxation rate. D) Percent muscle fatigue rate normalized to peak force at 100Hz. E) Area under the fatigue curve as a metric of overall skeletal muscle fatigue throughout the fatigue test. KD show significantly lower fatigue AUC compared to controls (*p=0.0359). All data presented are mean ± SD.

### iSOD2 KD muscle mitochondria display characteristics of impaired metabolic flexibility

Following four weeks of recovery from muscle function testing animals were euthanized and tissues were collected for High Resolution Respirometry (HRR), enzyme activity, and protein content via western blotting. For these HRR measurements, protocol 1 was used (Table 1). There were no significant sex differences in mitochondrial respiration in either KD or controls, indicating the SOD2 KD had similar effects on respiration between males and females. Since there were no sex differences, the sexes were combined for subsequent analyses. There were no differences between KD and controls under leak state with glutamate and malate. Complex 1 (CI) driven state 3 respiration with glutamate, malate, and pyruvate (GMP) is significantly lower in KD compared to controls (Figure 6A, p<0.0001), but there was no difference with just glutamate and malate. Maximal OxPhos capacity is also significantly lower in KD compared to controls (Figure 6A, p<0.0001). There were no significant differences in complex II or complex IV respiration between KD and controls. Taken together, these results corroborate our findings from the time course experiments. The decline in pyruvate driven respiration is consistent with previous reports on constitutive skeletal muscle SOD2 knockouts and is a hallmark of metabolic inflexibility (Muoio, Noland et al. 2012, Ahn, Ranjit et al. 2019, Koves, Zhang et al. 2023, Siripoksup, Cao et al. 2024).

**Figure 6.**
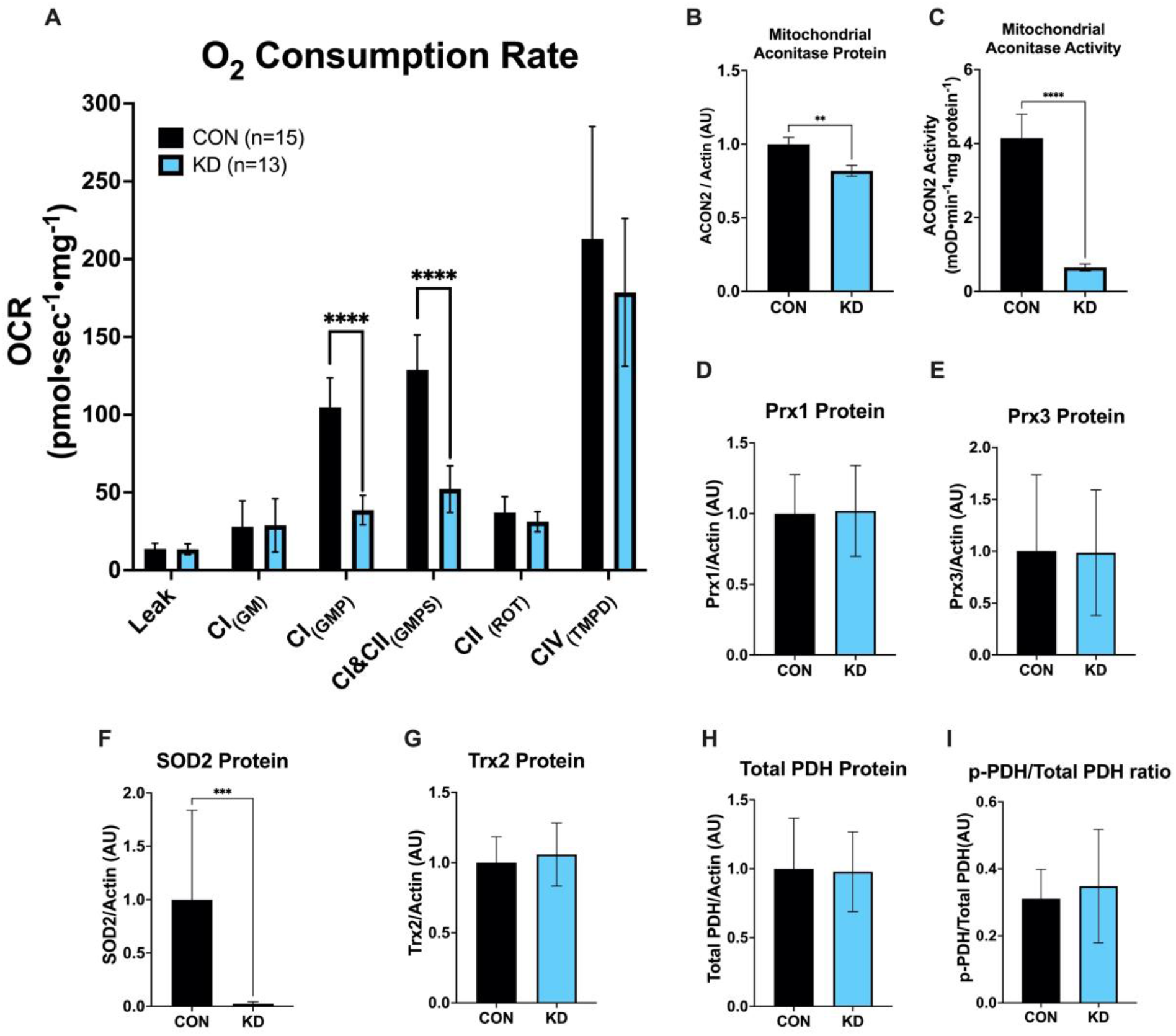
Oxygen consumption rate, enzyme activity, and protein content of select antioxidant and metabolic enzymes after 12 weeks of DOX administration. A) A standard SUIT protocol (Protocol 1) was used to assess mitochondrial function in permeabilized red gastrocnemius fibers. KD demonstrate significant impairments in pyruvate-driven respiration and maximal oxidative phosphorylation (****p<0.0001). B) Mitochondrial aconitase protein was significantly lower in KD compared to controls (**p<0.01). C) Mitochondrial aconitase enzyme activity was also significantly lower in KD compared to controls (****p<0.0001). D) Prx1 Protein content was no different between KD and CON. E) Prx3 protein content was no different between KD and CON. F) SOD2 protein content was significantly lower in KD (***p<0.001). G) Trx2 protein content was not different between KD and CON. H&I) Total PDH, and phospho-PDH were no different between KD and CON. Statistical tests are independent sample t-test.

We measured mitochondrial aconitase activity from isolated mitochondria as an indicator of mitochondrial oxidative stress (Murphy, Bayir et al. 2022). We also measured aconitase protein levels via western blotting to control for any changes in aconitase protein content that would affect the aconitase activity results. The KD group had lower ACON2 protein levels (Figure 6B, p<0.01) and lower aconitase activity compared to controls (Figure 6C, p<0.0001), similar to the results found in our time course experiments (Figure 2F), however this only partially explained the loss of ACON2 activity (Supplementary Figure S5) suggesting the remaining ACON2 enzyme was impaired by inactivation of ACON2 in KD muscle. There was almost complete ablation of SOD2 protein, consistent with our time course findings (Figure 6F, p<0.001). However, there were no differences in antioxidant proteins Prx1, Prx3, or Trx2 and no differences in total or phosphorylated pyruvate dehydrogenase (p-PDH/total PDH, Figure 6D-E, G-I). Representative images of western blot data presented in Figure 6 can be found in supplementary figure 6.

### iSOD2 KD is a model of impaired metabolic flexibility dependent on substrate order

Since we observed impaired respiratory capacity in permeabilized muscle fibers of iSOD2 KD animals with the addition of pyruvate, we wanted to test whether this impairment was also seen with other complex I substrates. To test this, we performed another series of HRR experiments in red gastrocnemius fibers. We split fibers from each animal into 3 different chambers to run respirometry experiments in parallel. We used protocols 2-4 for these experiments (Table 1). Leak respiration was no different between KD or controls for any protocol (Figure 7A-D). State 3 respiration with palmitoyl carnitine and beta hydroxybutyrate were similar to glutamate (Figure 7 A, C, & D – Protocol 1, 3, and 4, respectively). In SUIT protocol 3 and 4, the addition of pyruvate (Step 4) increased respiration significantly in the controls, but not in the KD muscle (Figure 7C-D). This impairment is almost identical to the impairment in protocol 1 when pyruvate is added after glutamate and malate. This impairment is not seen in protocol 2 when pyruvate was added before glutamate and palmitoyl carnitine (Figure 7B). Respiration from all 4 protocols with all complex 1 substrates are graphed in Figure 7E for easier visual comparison across protocols. In protocol 1, 3, and 4 there are significant impairments in KD muscle but not in protocol 2 (Figure 7E). These results demonstrate that pyruvate driven respiration is not impaired in skeletal muscle of iSOD2 KD when pyruvate is present before other substrates, but significantly impaired when other substrates are present before pyruvate, suggesting that there is some substrate level regulatory feedback mechanism controlling pyruvate oxidation. This control is dependent on the presence of other substrates that contribute NADH to complex I (glutamate, palmitoyl carnitine, or beta hydroxybutyrate). There were no significant differences in citrate synthase activity (Figure 7F), Complex I-V protein content (Figure 7G) but KD did show slightly lower CII enzyme activity compared to controls (Figure 7H, p<0.05).

**Figure 7.**
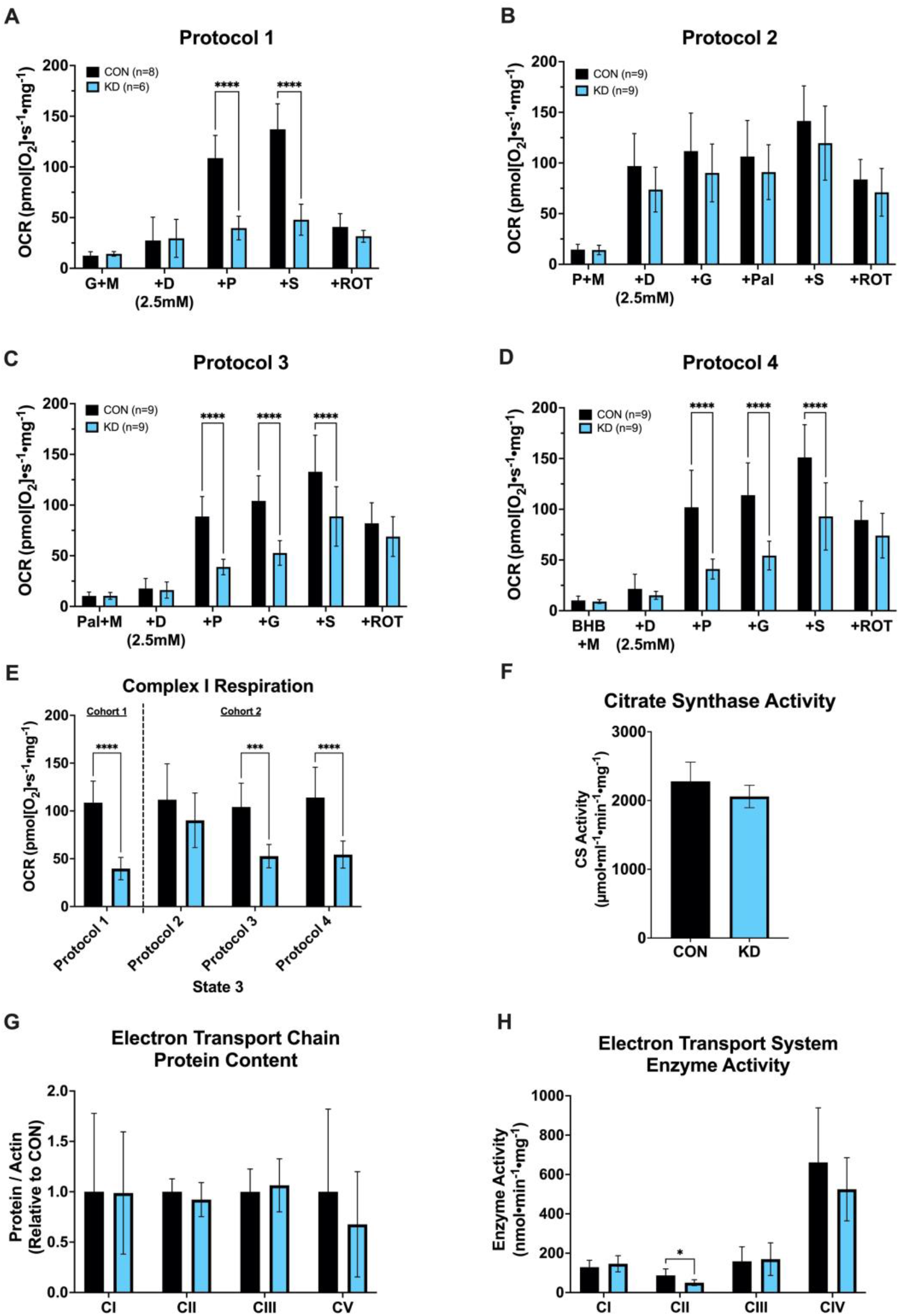
Substrate order significantly affects maximal oxygen consumption rate. A second cohort of female mice were treated with DOX continuously for 12 weeks followed by high resolution respirometry in permeabilized red gastrocnemius fibers. Muscle fibers were split into different chambers and parallel SUIT protocols were run simultaneously (Protocol 1-4). Substrate concentrations and order of addition into the chambers are illustrated in Table 1. A) Protocol 1 – adding glutamate and malate then pyruvate results in significant differences in respiration between KD and CON. These are results from females in Cohort 1 experiments and are graphed for illustrative purposes. B) Protocol 2 – No significant differences between controls and KD when pyruvate and malate are added first followed by glutamate (+G) and palmitate (+Pal). C) Protocol 3 – A significant impairment in respiration is seen when adding palmitate and malate before adding pyruvate, similar to the effects seen from Protocol 1. D) Protocol 4 – Again there is a significant impairment in respiration between KD and controls when adding beta hydroxybutyrate (BHB) and malate before adding pyruvate like Protocols 1 and 3. E) Complex 1-dependent respiration data pulled from graphs A-D for visualization purposes. For each protocol the bars graphed show all complex I substrates on board prior to the addition of succinate (Table 1 Step 4 for Protocol 1, and Table 1 Step 5 for protocols 2-4). F) No significant differences between KD and controls for citrate synthase activity. G) No significant differences between KD and controls for Complex I, II, III, and V protein content. H) There was a significant difference in Complex II enzyme activity between KD and controls but no other differences between KD and controls. All data presented are mean ± SD. Independent sample t-tests *p<0.05, **p<0.01, ***p<0.001, ****p<0.0001.

### Impaired pyruvate respiration in iSOD2 KD muscle is partially rescued with PDK inhibitor DCA but not rescued with thiol reducing agent DTT or superoxide scavenger MitoTEMPO

To understand the mechanisms underlying this acute regulatory inhibition of pyruvate oxidation we treated a third cohort of animals with DOX for 12 weeks, followed by tissue collection and permeabilization of red gastrocnemius fibers. We used HRR with the first 4 steps of SUIT protocol 1 and 2 which only vary by switching the order of glutamate and pyruvate (Table 1). Within each protocol permeabilized fibers were split into three different treatment conditions in the respiration buffer. Either 5mM dichloroacetate (DCA) a PDK inhibitor which would increase activity of PDH, 5µm UK5099, a mitochondrial pyruvate carrier (MPC) inhibitor as a negative control, or DMSO (Vehicle Control). Under vehicle conditions in protocol 1 there was a similar impairment in pyruvate driven respiration (Figure 8A, purple bars p<0.05). DCA treatment demonstrated a relative increase in pyruvate driven respiration in protocol 1 (Figure 8A, pink bars DCA effect, p<0.05), however glutamate driven respiration was abnormally low compared to vehicle, suggesting some inhibitory effect of DCA on glutamate respiration.

**Figure 8.**
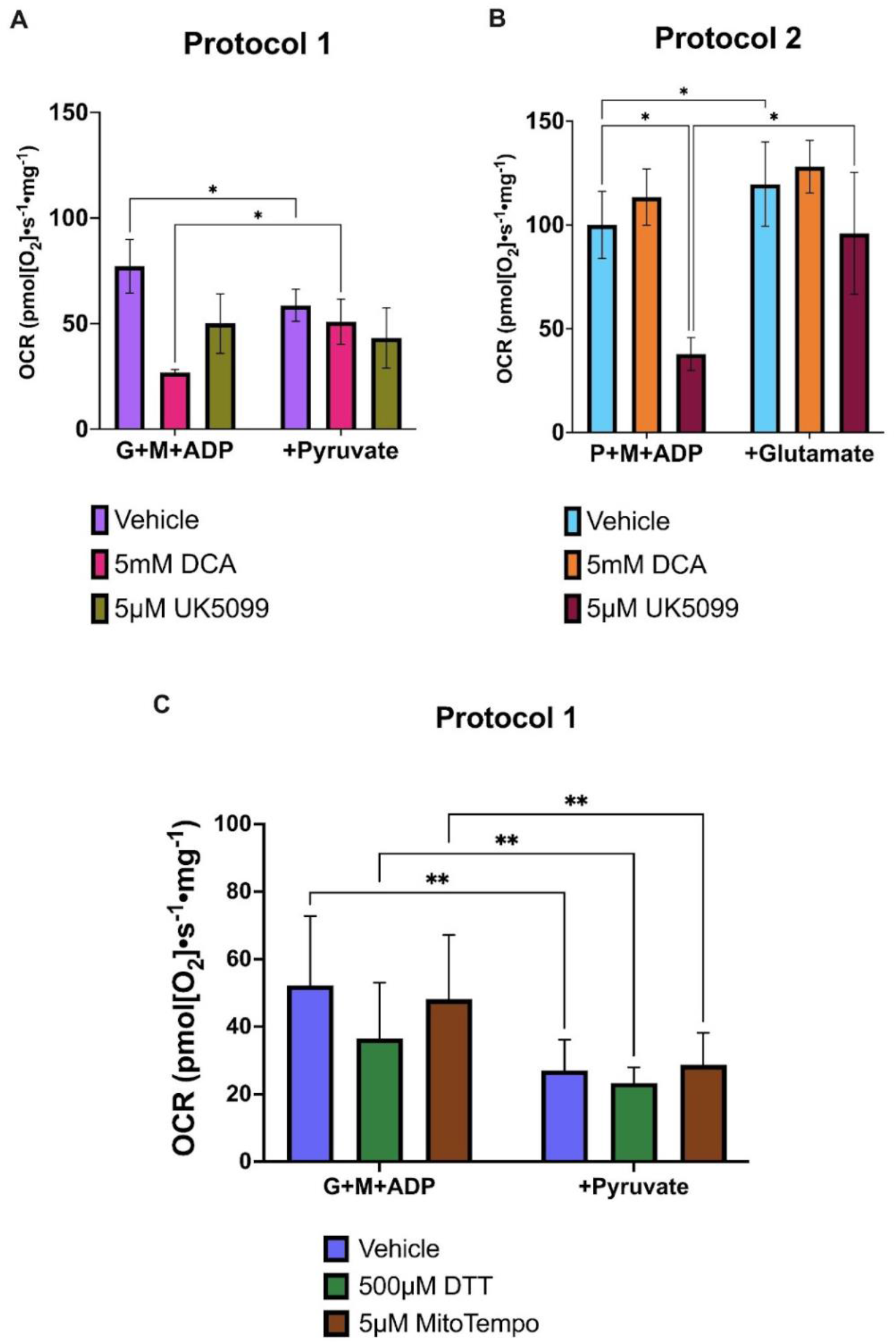
Allosteric regulation via redox or PDK phosphorylation dependent mechanisms do not explain impaired pyruvate driven respiration in permeabilized muscle fibers. A third cohort of animals were treated with doxycycline continuously for 12 weeks followed by tissue collection, gastrocnemius permeabilization and high resolution respirometry in iSOD2 KD animals only. Protocol 1 (Panel A) and Protocol 2 (Panel B) were used with *ex vivo* treatment of either Dichloroacetate (DCA), a Pyruvate Dehydrogenase Kinase (PDK) inhibitor or UK5099, a mitochondrial pyruvate carrier (MPC) inhibitor. A) Pyruvate driven impairment was seen in protocol 1 similar to previous experiments (Vehicle condition). Although there was a significant increase in pyruvate respiration with DCA treatment, the initial levels of respiration were low indicating some negative effect of DCA on glutamate driven respiration. UK5099 inhibited respiration under both substrate conditions. B) There were no impairments in pyruvate driven respiration when pyruvate was added first – similar to our previous experiments. DCA increased respiration but not significantly, while UK5099 inhibited respiration through decreased pyruvate transport into the mitochondrial matrix. C) Treating with DTT, a thiol reductant, or MitoTEMPO, a superoxide dismutase 2 mimetic, did not rescue the impaired pyruvate respiration (Two-way ANOVA with Tukey’s post hoc comparisons *p<0.05, **p<0.01)

When pyruvate is added first in iSOD2 KD muscle there is a normal increase in respiration and a slight additive effect of glutamate (Figure 8B, vehicle condition, p<0.05). DCA treatment did not have a significant effect on pyruvate driven respiration in protocol 2. UK5099 caused inhibition of pyruvate driven respiration in protocol 2 similar to the impaired pyruvate respiration in the vehicle condition in protocol 1. We also tested whether direct oxidation of protein thiols could explain this decline in pyruvate driven respiration. To do this we treated fibers with DTT (a thiol reductant) or MitoTEMPO (an SOD mimetic) in the respiration media followed by SUIT protocol 1. Neither DTT nor MitoTEMPO rescued the impaired pyruvate driven respiration (Figure 8C).

## Discussion

We describe a new model of mitochondrial oxidative distress in skeletal muscle using a Tetracycline Response Element (TRE) promoter to control short hairpin RNA targeted to SOD2 mRNA. The strength of this model over other constitutive knockout or inducible knockout models is that it is reversible, allowing researchers to test whether knockdown is necessary and sufficient for changes in metabolism and redox signaling or to test adaptive responses to mitochondrial oxidative distress after recovery from knockdown. The goal of this project was to characterize the time course of SOD2 knockdown and the resulting mitochondrial phenotype.

Results from this model indicate a delay between the elevation in matrix mitochondrial redox stress and the metabolic phenotype. SOD2 protein was almost completely absent by week 6 but the declines in mitochondrial energetics were not present until week 12 of recovery indicating that the effects of redox stress are not due to acute signaling responses such as post-translational modification of proteins, but rather require accumulated exposure to redox stress. Previous work has established that elevated mitochondrial oxidative stress in skeletal muscle drives decline in mitochondrial energetics and muscle function (Ahn, Ranjit et al. 2019, Zhuang, Yang et al. 2021). One of the findings from this model was an impaired pyruvate driven respiration in the KD following the addition of glutamate and malate. Our results indicate that this is among the first phenotypic changes to occur in response to mitochondrial oxidative distress. Importantly, the declines in pyruvate metabolism were reversed with SOD2 protein by week 24 of recovery. We can conclude from these time course results that the metabolic inflexibility is caused by mitochondrial oxidative distress.

While we observed robust declines in mitochondrial respiration, ROS production per unit of oxygen consumption, and aconitase activity, there were no major changes in gene expression responses in antioxidant defenses. We speculate that this is because superoxide, a negatively charged free radical, does not readily permeate the mitochondrial double membrane to drive gene expression changes in the nuclear compartment. Previous work in C2C12 cells demonstrate that mitochondrial retrograde redox signaling is driven by changes in mitochondrial thiol oxidation status and not superoxide production per se (Cvetko, Caldwell et al. 2021). It is also possible that the knockdown of SOD2 was not long enough to induce redox stress response signaling. Previous work using an inducible SOD2 knockout model also did not observe any significant changes in 4HNE (Zhuang, Yang et al. 2021), a maker of oxidative distress, indicating this model produces a relatively mild phenotype absent of changes in traditional markers of oxidative distress which is consistent with our lack of gene expression responses. This also highlights the importance of using appropriate markers of oxidative distress like aconitase activity which is more sensitive and specific to elevations in matrix superoxide levels than general markers of oxidative distress like 4HNE. Taken together, these time course results suggest the metabolic inflexibility is an early marker of elevated steady state levels of mitochondrial superoxide.

The iSOD2 KD model was originally developed and characterized by Cox et al., (Cox, McKay et al. 2018). In these experiments, animals were treated with DOX in utero for 4-8 days to knockdown SOD2. After 4 weeks of recovery from DOX administration, SOD2 protein was fully recovered in mouse livers. There were a host of biochemical and metabolic adaptations related to mitochondrial biogenesis and antioxidant defense system signaling, indicating a robust mitohormetic response in mouse liver tissue. Given those results, we expected the recovery dynamics in our model to be much faster than we observed. While we do not have an explanation for the significant delay in SOD2 recovery dynamics, it is possible the rtTA transcription factor remains activated for a sustained period of time following DOX administration. Future experiments will test this hypothesis.

### Skeletal Muscle Function and Fatigue

Skeletal muscle force production and calcium handling can be affected by oxidative distress (Andrade, Reid et al. 1998, Andersson, Betzenhauser et al. 2011, Baumann, Kwak et al. 2016). In aged muscle this is thought to stem, at least in part from modification of the ryanodine receptor and SERCA by elevated mitochondrial ROS production (Andersson, Betzenhauser et al. 2011, Umanskaya, Santulli et al. 2014). In these experiments, we did not observe differences in maximal isometric muscle force or contractile kinetics between iSOD2 KD and control muscle, however, there was a small but significant decrease in muscle fatigue resistance in the iSOD2 KD muscle assessed by fatiguing muscle contractions. While this effect is relatively small in this high intensity isometric contraction protocol the effects of this decreased muscle endurance and recovery may become more functionally important under more sustained protocol, with repeated bouts of contraction or after a longer period of SOD2 KD.

### Mitochondrial Substrate Utilization and Control

Skeletal muscle is a critical tissue for whole body metabolic health and is a major site of glucose and fatty acid disposal. Skeletal muscle mitochondria play a major role in metabolic control, adjusting substrate utilization during periods of fasting, feeding, and energetically demanding conditions such as exercise to maintain metabolic and redox homeostasis. Metabolic flexibility, defined as the ability to respond to changes in metabolic demand (Goodpaster and Sparks 2017), is thought to be controlled at least in part by skeletal muscle mitochondria (Muoio, Noland et al. 2012, Muoio 2014). In healthy mitochondria adding multiple substrates such as glutamate and pyruvate under state 3 conditions has additive effects on oxygen consumption. Additive effects on oxygen consumption have been discussed in the context of complex I and complex II substrates previously, which are relevant for insulin resistance and type 2 diabetes (Boushel, Gnaiger et al. 2007, Gnaiger 2009). Others have described impaired pyruvate respiration after the addition of palmitoyl carnitine due to incomplete fatty acid oxidation, but there were no inhibitory effects when other, non-fatty acid, substrates were added first (Koves, Zhang et al. 2023). To our knowledge this study is the first to report that the presence of multiple substrates, glutamate, beta-hydroxybutyrate, or palmitoyl carnitine, entering the ETS through multiple pathways, inhibit pyruvate oxidation and thereby limit mitochondrial metabolic flexibility. These results suggest that under *in vivo* conditions with a complex mixture of substrates, elevated matrix redox stress may contribute to reduced metabolic flexibility and reduced ability to use glucose to drive oxidative phosphorylation.

Pyruvate oxidation is allosterically controlled by phosphorylation/dephosphorylation of pyruvate dehydrogenase complex (PDHC) by pyruvate dehydrogenase kinase (PDK) and pyruvate dehydrogenase phosphatase (PDP), respectively. PDK phosphorylation of PDHC decreases PDHC activity, so hyperactivation of PDK and phosphorylation of PDHC is a potential mechanism underlying inhibited pyruvate oxidation in the SOD2 KD model. Treatment with dichloroacetate (DCA), a pharmacological inhibitor of PDK, partially rescued the inhibition of pyruvate oxidation in the presence of glutamate in the iSOD2 model, suggesting that the impaired pyruvate oxidation is at least partly controlled by PDHC phosphorylation.

Previous work has shown that pyruvate oxidation is also controlled by glutathionylation of the mitochondrial pyruvate carrier (MPC). Experimentally increasing MPC s-glutathionylation inhibits pyruvate uptake and oxidation within the mitochondria (Gill, O’Brien et al. 2018). In the current experiments, treatment with the MPC inhibitor UK5099 mimicked the impaired pyruvate respiration. However increased protein s-glutathionylation of MPC was not the mechanism responsible for impaired pyruvate respiration in iSOD2 KD because treatment with DTT did not rescue this effect. Additionally, mitoTEMPO, a superoxide dismutase mimetic, did not rescue pyruvate driven respiration. These results suggest a potential buildup of intermediate metabolites that cause feedback inhibition of PDH through PDK activation and the PDK activation is independent of oxidative post translational modifications, although further research is needed to determine what metabolite may be activating PDK and if this is the sole mechanism driving impaired metabolic flexibility.

One proposed mechanism that leads to metabolic inflexibility in skeletal muscle is the incomplete oxidation of fatty acids (Koves, Ussher et al. 2008). While we did not measure metabolic intermediates after mitochondrial respiration experiments, it is possible the presence of multiple mitochondrial substrates in iSOD2 KD muscle causes buildup of TCA cycle intermediates and the major control mechanism to minimize this build up is to shut down pyruvate oxidation. Indeed, impaired pyruvate metabolism occurs under several other experimental conditions in skeletal muscle. The addition of palmitoyl carnitine to mitochondria respiring with pyruvate and malate causes a decrease in efficiency and proton conductance in wild type muscle mitochondria (Koves, Zhang et al. 2023). These inefficiencies are due to bottle necks created by incomplete fatty acid oxidation and limited availability of free carnitine and free CoA (Koves, Zhang et al. 2023). Others have shown impaired pyruvate metabolism in sedentary mouse models related to decreased phosphatidylethanolamine (PE) abundance in the mitochondrial membrane (Siripoksup, Cao et al. 2024). Despite different control mechanisms between these models, impaired pyruvate metabolism appears to be a consistent feature. Future research is required to determine the mechanism of impaired pyruvate oxidation in the iSOD2 KD model and how this relates to other models of metabolic dysfunction.

### Conclusions

This project has provided a characterization of a new inducible and reversible SOD2 knockdown model. The main phenotype of the SOD2 KD is the inhibition of mitochondrial pyruvate oxidation, associated with elevated matrix oxidative distress, that is reversed as SOD2 protein expression levels recover following reversal of the inhibition. These results demonstrate the power of this model for investigating the relationship between mitochondrial redox biology and metabolic inflexibility associated with conditions such as diabetes and obesity. This has important implications for the interpretation of previous respirometry experiments because impaired pyruvate respiration is only observed when other substrates are present. In some previous studies there were no differences in respiration between diabetic and control skeletal muscle, however these experiments did not use multiple substrates to assess additive or order effects (Boushel, Gnaiger et al. 2007, Phielix, Schrauwen-Hinderling et al. 2008). It is possible these experiments would see impaired pyruvate respiration in diabetic or insulin resistant patients if pyruvate was added after other complex I substrates. Future experiments are needed to understand the mechanisms behind the impaired pyruvate oxidation, but these results are consistent with previous studies on impaired metabolic flexibility in skeletal muscle. This iSOD2 KD model will provide a useful tool to the field of redox biology and skeletal muscle metabolism in future studies.

## Acknowledgements

We would like to thank Dr. Gerald Shadel for his help troubleshooting the development of the iSOD2 KD mice. We would like to thank Dr. John McCarthy for generously donating the HAS-rtTA mice. We would like to thank Dr. Neal Paragas for access and assistance with the IVIS Spectrum *In Vivo* Imaging System. We would like to thank Dr. Anna Scott for performing the electron transport system complex activity assays.

## Funding Sources

R21 AG083241 to DJM

R01 AG078279 to DJM

F32 AG074655 to

ELO P30 AT074990 to DJM

All figures created in Biorender

---

**Supplementary Figure S1.**
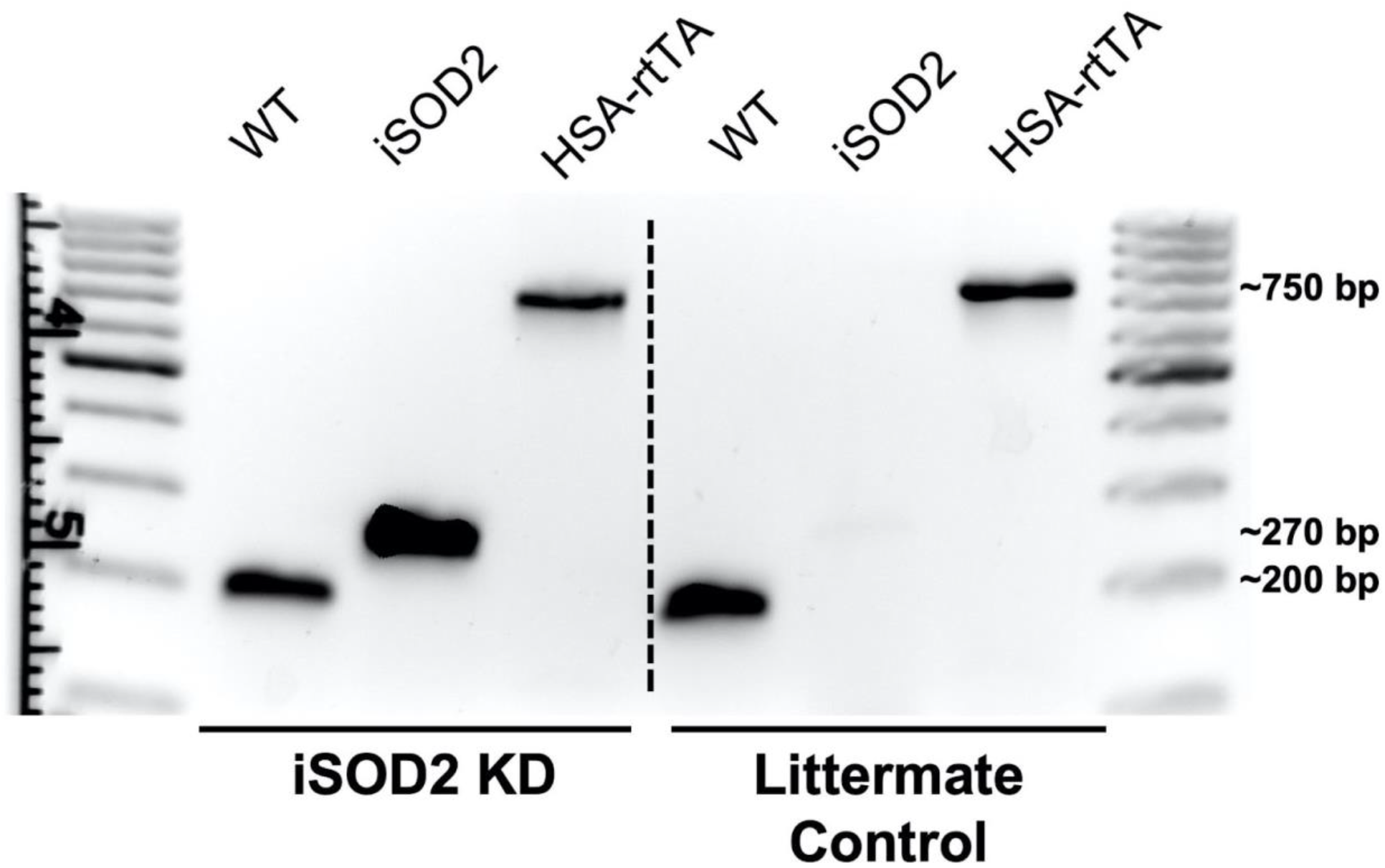
Representative image of genotype screening for an iSOD2 KD animal and a littermate control. WT – wild type allele migrates to approximately 200 bp, iSOD2 – inducible SOD2 KD allele migrates to approximately 270bp, HSA-rtTA – Human skeletal actin driven reverse tetracycline transactivator allele migrates to approximately 750bp. To induce SOD2 knockdown with doxycycline both the iSOD2 and the HSA-rtTA alleles need to be present. Littermate controls need to have only the HSA-rtTA allele to be the Doxycycline fed controls.

**Supplementary Figure S2.**
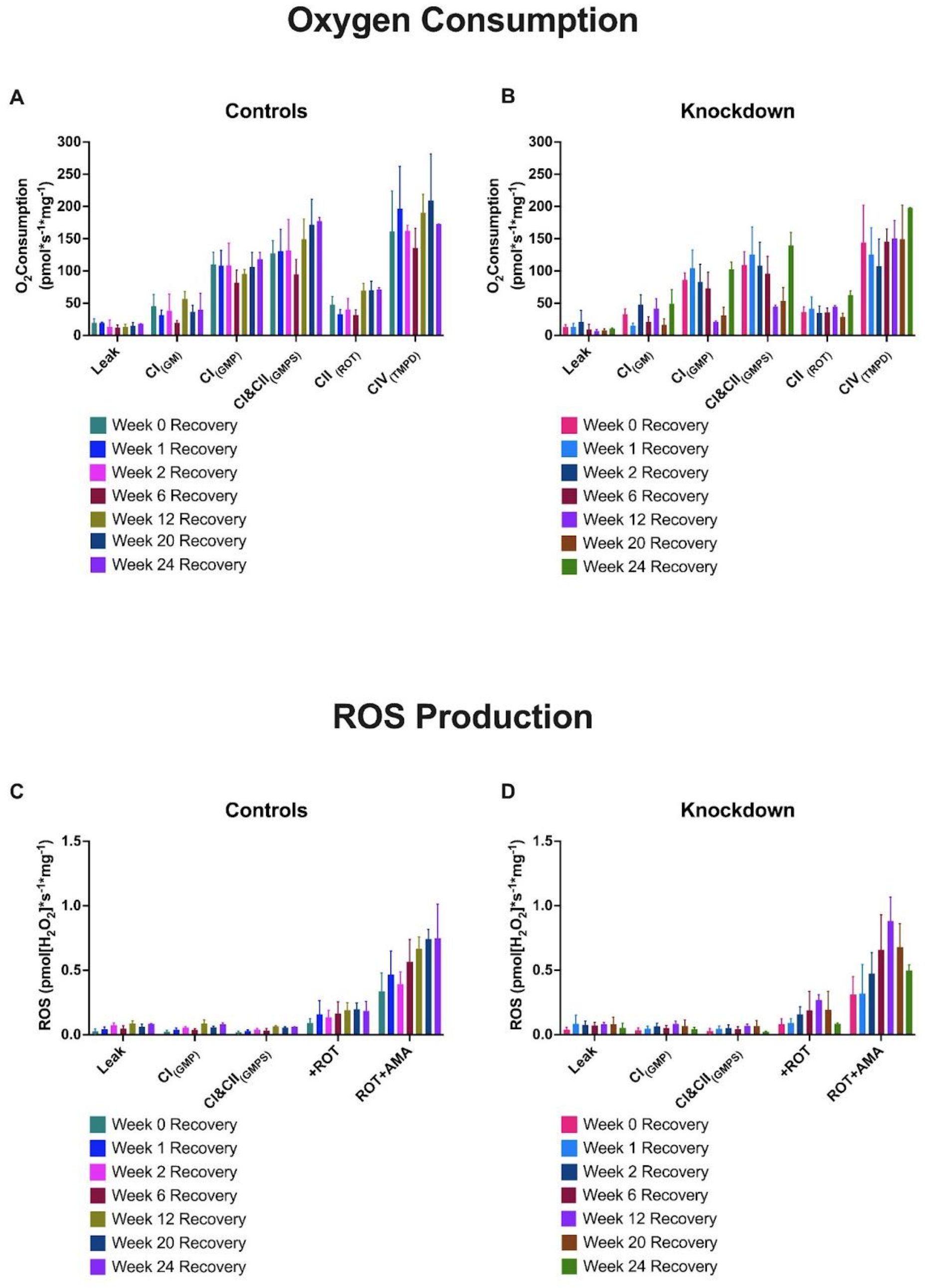
Mitochondrial oxygen consumption and ROS production rates in permeabilized red gastrocnemius muscle fibers at various recovery time points after 3-week DOX administration. Protocol 1 from table 1 was used for these experiments. Leak, Complex I state 3 respiration, maximal oxidative phosphorylation, Complex II driven respiration, and complex IV respiration for controls and KD (Panel A&B). ROS production rate during leak and state 3 respiration, and rates of ROS production during Reverse Electron Transport (RET) from Rotenone inhibition of Complex I, and complex III driven ROS production from Antimycin A treatment (Panel C&D). When parsed by sex, there were no significant differences between male and female controls, or within sex effects of genotype for any substrate While there were no significant differences in ROS production rates between KD and controls, there is a pattern of increased ROS production in both Rotenone and Antimycin A treatments by week 12 of recovery in the KD animals which corresponds to the lowest OCR measured in all experiments. All data presented are mean ± SD.

**Supplementary Figure S3.**
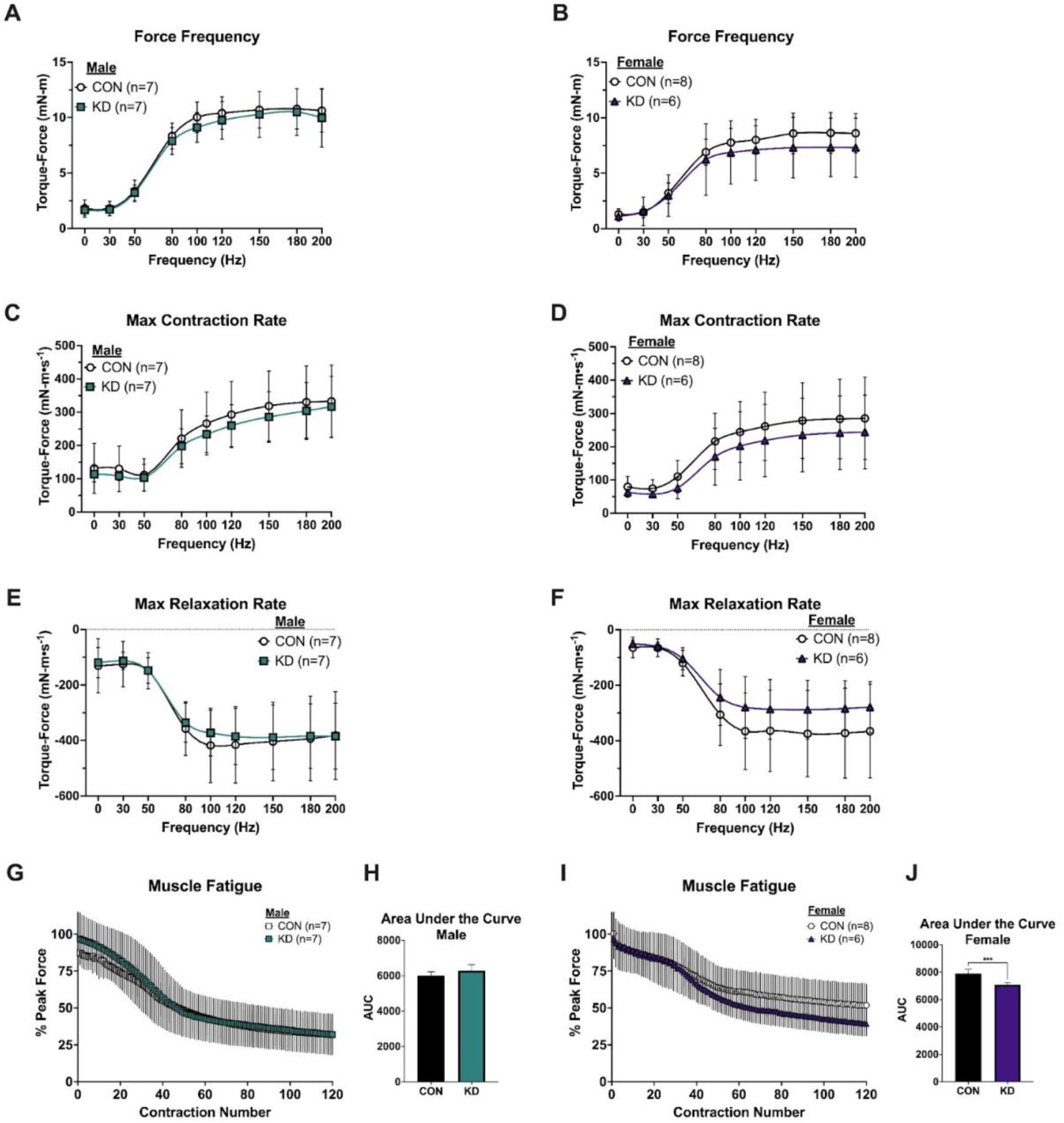
*In vivo* skeletal muscle function and fatigue following 8 weeks of continuous DOX administration parsed by sex. Force Frequency curve for male (A) and female (B) cohorts. Maximal contraction rate of male (C) and female (D) cohorts. Maximal relaxation rate for male (E) and female (F) cohorts. Muscle fatigue (G) expressed as a percentage of peak force and area under the curve (H) for males. Muscle fatigue (I) expressed as a percentage of peak force and area under the curve (J) for females. KD females show significantly lower fatigue AUC compared to controls (***p=0.0002). All data presented are mean ± SD.

**Supplementary Figure S4.**
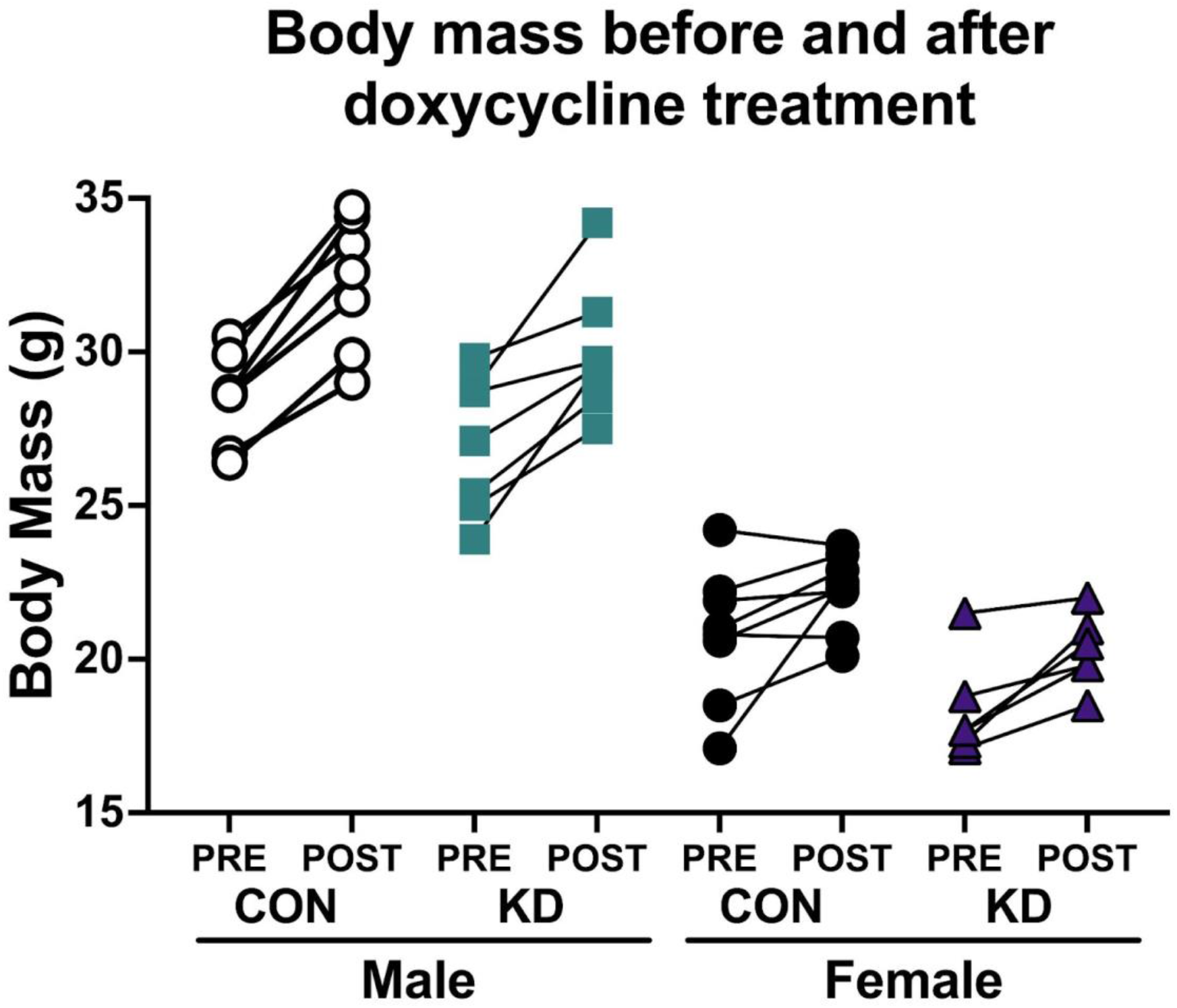
Body mass before and after 12 weeks of continuous DOX treatment in male and female KD and CON animals. All groups increased body mass, which can be explained by the normal maturation process of mice from 3-6 months of age.

**Supplementary Figure S5.**
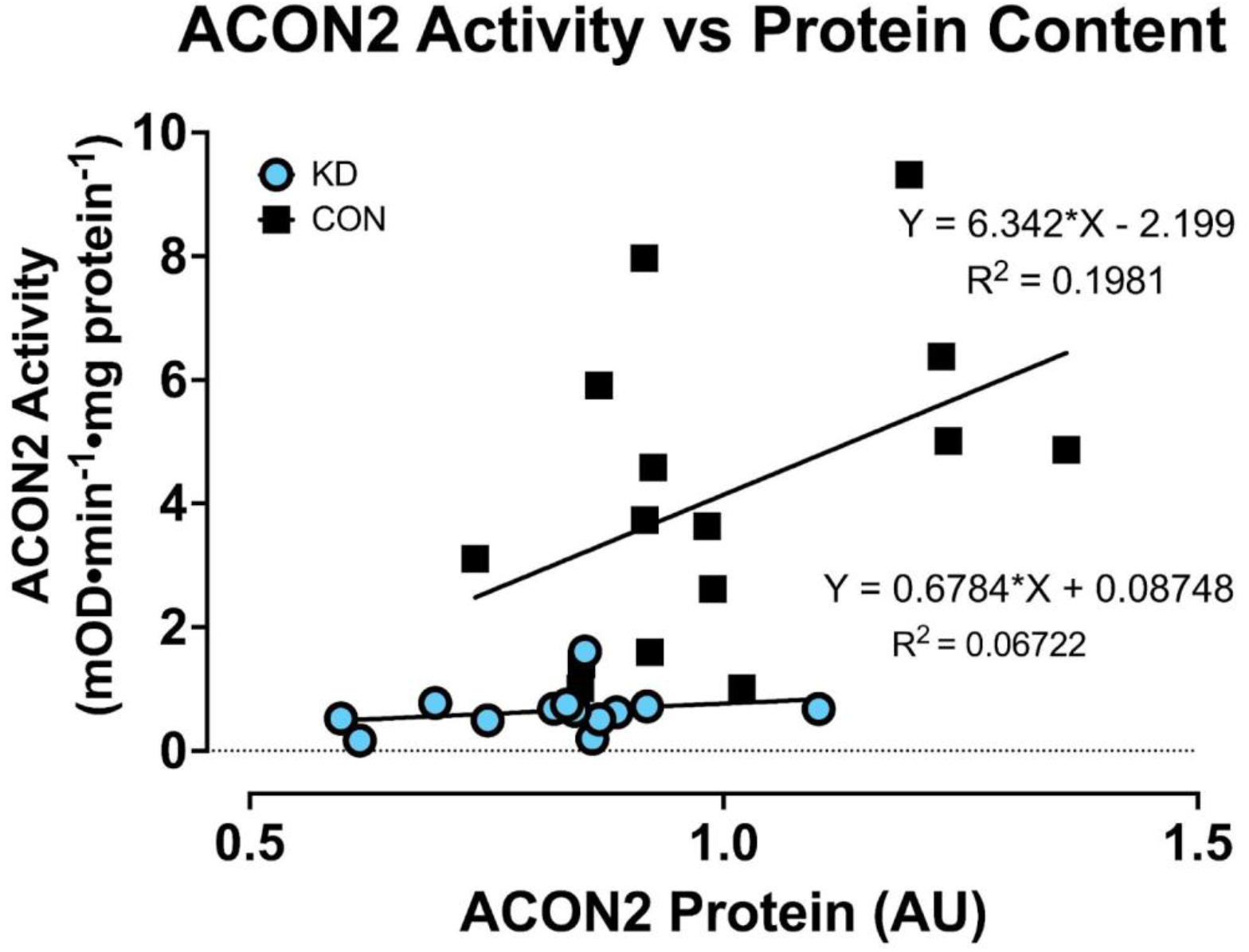
Simple linear regression of ACON2 protein content versus ACON2 activity in CON and KD muscle. Despite there being protein present in KD samples, there is minimal enzyme activity indicating an oxidative stress insult to iSOD2 KD mitochondria.

**Supplementary Figure S6.**
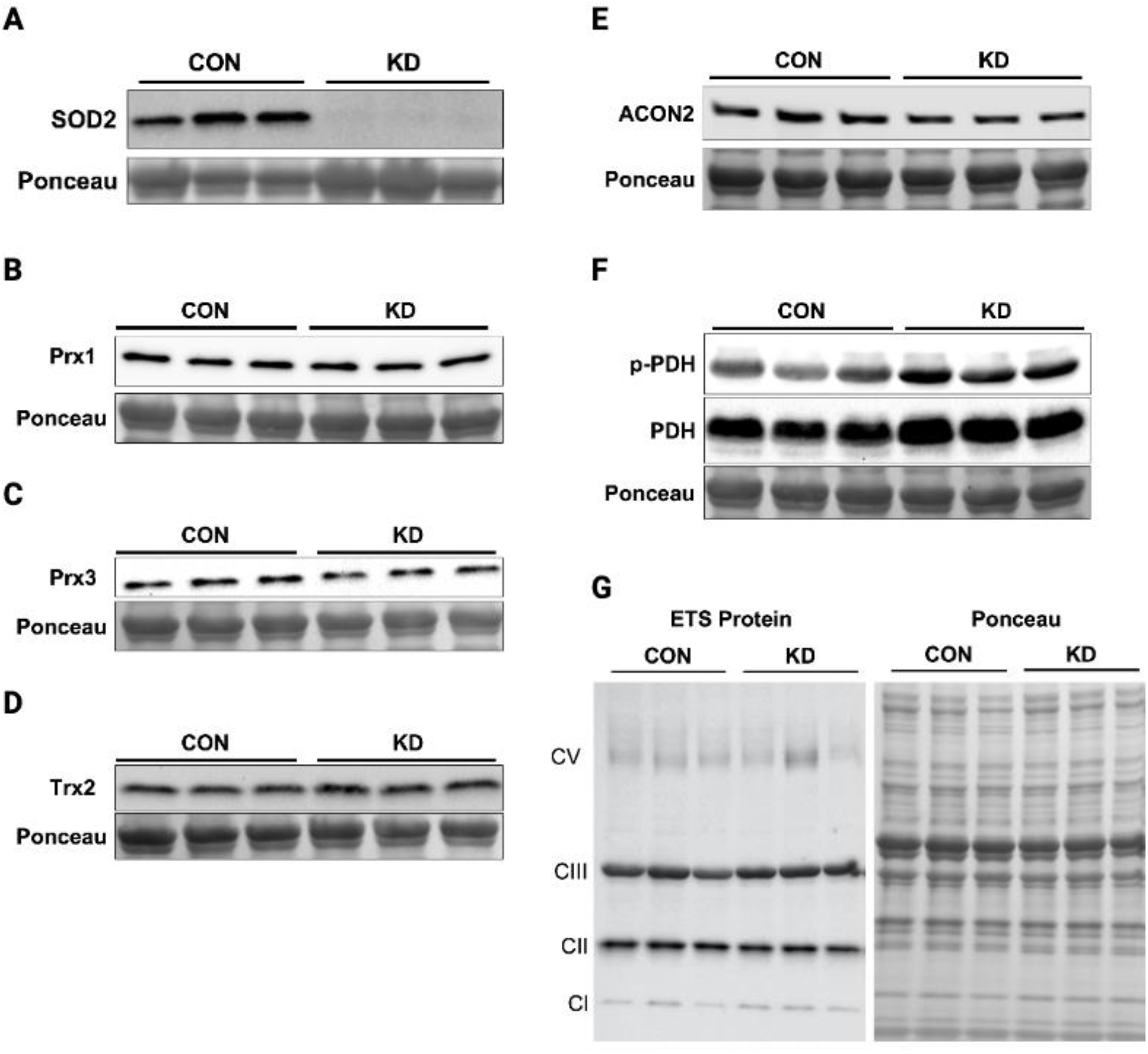
Representative images of western blots in gastrocnemius muscles treated with Doxycycline for 12 weeks. A) SOD2, B) Prx1, C) Prx3, D, Trx2, E) ACON2, F) phospho-PDH and total PDH, corresponding to data shown in Figure 6. G) ETS proteins from Complex I, II, III, and V corresponding to data shown in Figure 7. Complex IV protein was not able to be identified in either group. All proteins were normalized to total protein loaded per well using a Ponceau S stain.

**Supplementary Table 1.** TABLE S1. ANTIBODY LIST.

| ANTIBODY | Company | Catalog Number | Species | Dilution | Primary Ab Buffer Recipe |
| --- | --- | --- | --- | --- | --- |
| SOD2 | Abcam | Ab13534 | Rabbit | 1:1000 | 1% NFDM in 0.1% TTBS |
| ACON2 | Abcam | Ab129105 | Rabbit | 1:1000 | 5% NFDM in 0.1% TTBS |
| PRX1 | Abcam | Ab109498 | Rabbit | 1:1000 | 5% NFDM in 0.1% TTBS |
| PRX3 | Abcam | Ab73349 | Rabbit | 1:1000 | 5% NFDM in 0.1% TTBS |
| TRX2 | Abcam | Ab185544 | Rabbit | 1:5000 | 5% BSA in 0.1% TTBS |
| PDH | Cell Signaling Technology | 3205 | Rabbit | 1:1000 | 5% BSA in 0.1% TTBS |
| P-PDH | Cell Signaling Technology | 31866 | Rabbit | 1:1000 | 5% NFDM in 0.1% TTBS |
| OxPhos Cocktail | MitoSciences | 604 | Mouse | 1:1000 | 1% BSA in 0.1% TTBS |

**Supplementary Table 2.**
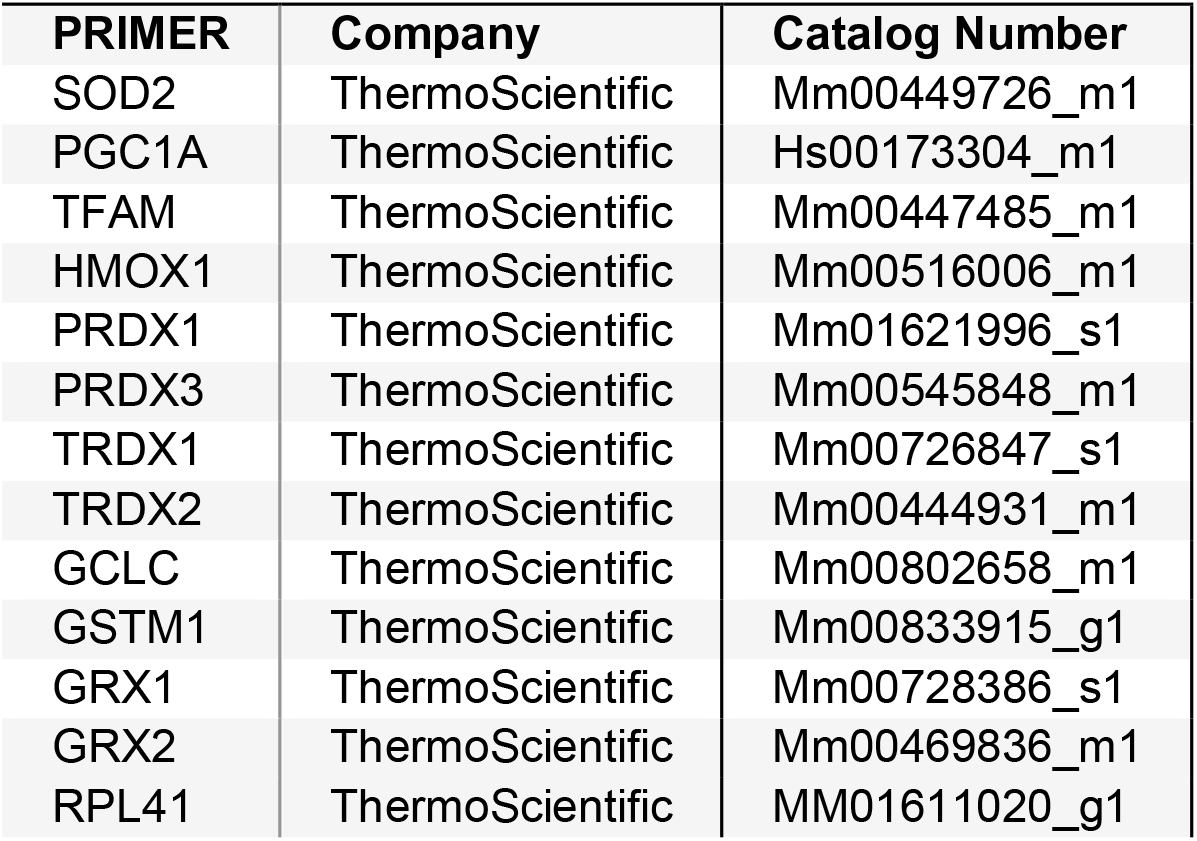
TABLE S3. RT-QPCR PRIMERS.

| PRIMER | Company | Catalog Number |
| --- | --- | --- |
| SOD2 | ThermoScientific | Mm00449726_m1 |
| PGC1A | ThermoScientific | Hs00173304_m1 |
| TFAM | ThermoScientific | Mm00447485_m1 |
| HMOX1 | ThermoScientific | Mm00516006_m1 |
| PRDX1 | ThermoScientific | Mm01621996_s1 |
| PRDX3 | ThermoScientific | Mm00545848_m1 |
| TRDX1 | ThermoScientific | Mm00726847_s1 |
| TRDX2 | ThermoScientific | Mm00444931_m1 |
| GCLC | ThermoScientific | Mm00802658_m1 |
| GSTM1 | ThermoScientific | Mm00833915_g1 |
| GRX1 | ThermoScientific | Mm00728386_s1 |
| GRX2 | ThermoScientific | Mm00469836_m1 |
| RPL41 | ThermoScientific | MM01611020_g1 |

